# Direct and indirect effects of the increase in atmospheric CO_2_ and temperature on groundwater organisms

**DOI:** 10.1101/2023.09.13.557665

**Authors:** Susanne I. Schmidt, Miroslava Svátková, Vít Kodeš, Tanja Shabarova

**Affiliations:** Biology Centre CAS, Institute of Hydrobiology, Na Sádkách 7, 370 05 České Budějovice, Czech Republic; Czech Hydrometeorological Institute, 143 00 Prague, Czech Republic

**Keywords:** groundwater ecosystem, weathering, fauna, prokaryotes, silica, Earth’s Critical Zone

## Abstract

Atmospherically rising temperature and CO_2_ impact all freshwater systems, including groundwater. Increasing CO_2_ leads to more intense weathering of silicate rocks. Here, we tested whether the increased levels, the weathering, or rather the increasing temperature, impacted on fauna and prokaryotes in the groundwater ecosystem. We conducted the analyses separately for deep, i.e. secluded, and shallow, quaternary aquifers which exchange with the surface more intensely. Organism abundances and relative composition did not correlate with temperature or CO_2_ levels. While many organisms rely on silica, in contrast, we found negative correlations between silica and fauna. The increases in silica over time, i.e. temporal trends, also partly correlated negatively with organisms. We hypothesize that the unexpected negative correlations are not direct effects, but indirectly indicate that groundwater communities do not adapt rapidly enough to changes. This jeopardizes future drinking water production which relies on the self-cleaning ecosystem services in groundwater.

## Introduction

Atmospheric CO_2_ levels and temperature have risen globally, particularly in the past 30 years, largely due to industrialization (IPCC, 2022) and/or anthropogenically increased CO_2_ (Penman et al., 2020). Consequently, by diffusion into water, aqueous CO_2_ concentrations have risen (Cole, 2013). Sea water is recognized to be one major sink for CO_2_ (Sabine et al., 2004), and in rivers, HCO_3_^-^ concentrations, most probably derived from increased CO_2_, have shown higher concentrations (Meybeck, 1993). However, it is much less clear whether groundwater is a sink or source for global CO_2_ levels (Boerner & Gates, 2015; Nydahl et al., 2020). More than likely, the net import of CO_2_ into groundwater has risen, via the concentrations that have increased in soil (Bakalowicz, 1994). The reason for the latter is that higher temperatures led to e.g. plant growth and in parallel increasing soil microbial activity and thus an increase in CO_2_ production through respiration (Conant et al., 2011). The increased CO_2_ and HCO_3_^-^ concentrations are subsequently recharged into groundwater. This increased CO_2_ import will immediately lead to increased acidity which, in turn, will dissolve minerals, and thus, fuel weathering (for a summary of CO_2_-driven weathering, see Supplementary Information SI 1). In fact, in parallel to sea water acting (still) as a CO_2_ sink, rock weathering under the consumption of CO_2_ is regarded as one mechanism stabilizing earth’s climate (Brantley et al., 2023; Li et al., 2016). Such geochemical processes are estimated to have absorbed the equivalent of about half of the carbon dioxide that was emitted each year in the 2012-2021 decade (Friedlingstein et al., 2022). While it is not even very clear which effects the increased CO_2_ levels have on freshwater organisms in general (Weiss et al., 2018), the effect on groundwater organisms which differ from surface organisms in many traits (Hose et al., 2022), is even less clear. More is known about temperature: while microbial numbers and diversity increased with temperature, faunal abundances and diversity decreased (Brielmann et al., 2009), and above 12°C, a fauna community shift in mid-European groundwater assemblages was observed (Spengler & Hahn, 2018; see also below).

Climate change in general threatens groundwater ecosystems, but we do not know yet, how which groundwater ecosystems will react, and which will be the driving impacts (Mammola et al., 2019). Thus, we do not know what will happen to the ecosystem functions that the ecosystems deliver, e.g. self-cleaning (Griebler & Avramov, 2015). A decline of groundwater ecosystem functions may threaten drinking water production which globally relies on groundwater to at least 50% (UNESCO World Water Assessment Programme, 2022).

Increased CO_2_ levels, in contrast to the clear direct effects from temperature, might have contradictory effects. They might provide the electron acceptor and/or carbon substrates for microbial reactions, such as the Calvin cycle, or the oxidation of reduced sulphur and nitrogen compounds, such as sulphide and ammonia (Herrmann et al., 2015; Schmidt et al., 2017). Primary production in groundwater, without light, is chemolithoautotrophic. Increased CO_2_ might boost this primary production. Most groundwater food webs are likely bottom-up controlled and thus rely on organic import via recharge, or, sometimes in parallel (Herrmann et al., 2020; Kellermann et al., 2012), on primary production. Groundwater food webs might thus benefit from increased CO_2_ import and one might expect increased prokaryote and consequently fauna biomasses. In contrast, surface water crustacea have been shown to be negatively affected by increased CO_2_ values (Hasler et al., 2017; Langer et al., 2017; Weiss et al., 2018). Fauna might thus not be able to make use of the increased microbial primary production, which would lead to higher-than-expected microbial biomass.

In parallel to the anticipated direct effects of groundwater temperature and the unclear direct effect of groundwater CO_2_ on fauna, indirect effects have to be taken into account. One major indirect effect of increased CO_2_ may be the increased silicate weathering (“The gradual removal of atmospheric CO_2_ through dissolution of silicate and carbonate rocks”; IPCC, 2022), and consequently, increased silica concentrations, and their potential impact on all organisms, not only fauna. Here, we use silica to mean generically, the dissolved element Si (silicon), and ionic forms, such as SiO_4_^2-^ etc., while we use silicate to indicate the undissolved rock mineral. Increased silica concentrations are not expected to harm organism communities. Silicon, Si, is the second-most abundant element on earth. If it was harmful in itself, we would expect a lot less organisms anywhere near silica rock outcrops. However, silica may be a (limiting) nutrient for organisms that rely on it, such as diatoms, foraminifera, and radiolaria (Lampert & Sommer, 2007; Moore et al., 2013). An increase in dissolved silica may therefore benefit organisms. Organisms, and thus the ecosystem functions they deliver, evolved alongside the silica weathering since geological times. However, it is unclear whether organisms stay unfazed during comparably rapid changes to the silica weathering patterns as have been evolving recently. Maher (2010) related extinction events to periods with major changes in rock weathering.

We tested the direct effects of current temperature, redox potential, CO_2_, HCO_3_^-^, and silica on prokaryote numbers which are at the bottom of the food web, and on fauna, i.e. organisms being held back by 70 µm mesh size net, which are largely at the top of the food web, in a set of Western Czech groundwater aquifers. We classed fauna into high level taxa and calculated biomasses from the abundances which we also correlated to the current physical and chemical values. The current values at the period of sampling may, however, be the result of changes that occurred in the past decades. We assume that where such changes occurred, we will observe trends in temperature, CO_2_, HCO_3_^-^, and silica over the past decades. Therefore, we used these trends as proxies for the magnitude of (climate) change in the aquifers. Current fauna assemblage structure might reflect the changes that occurred in the past decades, since fauna integrates over a longer time. We therefore related the trends of the past decades to current organism numbers, to check whether decade-long changes predicted current fauna assemblages better than nowadays physical and chemical conditions do. We also related the composition of the fauna assemblages to the current values and the trends.

Shallow aquifers are more vulnerable than deep, secluded ones, because they are in more intensive exchange with the surface and the atmosphere (Calmels et al., 2011). Effects from climate change are likely to trickle down only slowly from the surface into deeper realms. These deeper aquifers usually also receive less import from the surface and thus, support lower numbers of less diverse organisms (Schmidt & Hahn, 2012). Therefore, we separated analyses for deep and shallow aquifers, as characterized by the Czech Hydrometeorological Institute. In addition to these clear-cut groups, the redox potential may be seen as an indicator for the vulnerability, or the degree of exchange with the surface, of an aquifer: usually, electron acceptors such as oxygen, nitrate, and sulphate are reduced during groundwater recharge and along the groundwater flow path, mainly by prokaryotes, while electron donors may be released from minerals (Meckenstock et al., 2015). Thus, the redox potential decreases and can be used to confirm the grouping into shallow, high-exchange, and deep, low-exchange aquifer groups. With the redox potential being an indicator for import from the surface, we suspected less organisms at lower redox potential. Many of the chemolithoautotrophically active microorganisms use CO_2_ and thus, chemolithoautotrophic conditions may compete with the climate-change-induced CO_2_-driven silicate weathering.

Since redox reactions at lower redox potentials yield less free energy (Aquilina et al., 2023), we expected prokaryote numbers, and consequently, faunal numbers, to be lower at lower redox potentials. However, fauna is not only limited by resources, but also by oxygen. Some protozoa are anaerobic (Fenchel & Finlay, 2008) and avoid oxygen. In the current contribution, however, we focus on metazoa, i.e. multicellular organisms, and use the term “fauna” for them. Groundwater fauna is much better adapted to low oxygen values than surface fauna (Malard & Hervant, 1999), but 1 mg L^-1^ DO seems to be a limit below which the vast majority of metazoa cannot survive, unless they undergo tight symbioses, which was so far only observed in the deep ocean, not in groundwater (Danovaro et al., 2010).

Our hypotheses were the following: 1) over the past decades, the increases in temperature, CO_2_, HCO_3_^-^, and silica, were significant in the investigated wells, while redox potential and DO stayed unchanged; 2) the increases were higher in shallow aquifers which are more vulnerable and exchange more intensely with the atmosphere; 3) there were direct effects of current temperature, DO, CO_2_, HCO_3_^-^, redox potential, and silica on current organism assemblages, with higher numbers of prokaryotes with higher temperature, DO, CO_2_, HCO_3_^-^, redox potential, and silica, and lower numbers and biomasses of fauna with higher temperature, CO_2_, HCO_3_^-^, but higher numbers with higher levels of DO, redox potential, and silica; 4) the higher the respective absolute slope of the trend was over the past decades, the lower were nowadays organism numbers and fauna biomass; the reason for this is that the more distinct the trends were, the more rapid the change must have occurred this would have been detrimental to groundwater organisms which are not adapted to rapid changes, 5) current faunal assemblage structure reflects not only current levels, but also the trends of the past decades.

## Methods

A subset of the wells monitored by CHMI (Czech Hydrometeorological Institute) were sampled during the respective spring and autumn campaigns during the years 2019 to 2021 (for a partial, preliminary account, see Schmidt et al., 2021). The wells were chosen according to geographical proximity to other study sites and according to logistic restraints. The coordinates of the location of the wells are given in Table S1, and the position is indicated in the overview map Figure S1, in the Supplementary Information SI 2.

The chemical and physical results for those wells where fauna was sampled, were extracted from the CHMI database. See Supplementary Information SI 3 for the evaluation of CO_2_. Molar ratios of Na/Ca versus Mg/Ca were calculated to show which type of rock weathering might have dominated (Gaillardet et al., 1999; Négrel et al., 1993).

For the analyses of trend over time, the complete data set, covering the whole monitored period, was used, while for the analyses of the recent or current environment of the organisms, the chemical and physical values were averaged over the years 2019 – 2021. The recent chemical and physical conditions were correlated to faunal abundances, faunal biomasses, and prokaryote numbers, and linear regression parameters were calculated. The slopes of the linear trends of these variables over the past decades were also correlated to faunal abundances, biomass, prokaryote numbers, and relative assemblage composition (see below: “envfit”), to reflect whether recent developments had shaped organisms.

Fauna was sampled first with a 70 µm pore net (manufactured by Institut für Grundwasserökologie GmbH, Landau, Germany) lowered down the well and drawn up at arm’s length for ten times to scoop up bottom-living fauna (Bou, 1974; Matzke & Hahn, 2002; Schmidt et al., 2004). Subsequently, from the pumped water stream, 50 L aliquots of pumped water were taken and sieved through the same net. It was checked whether there was an obvious difference in numbers of fauna sampled by pumping or with the net draws, related to the volume that was estimated to have passed through the net (Further information on sampling is given in Supplementary Information SI 4). Fauna biomasses were estimated from abundances based on biomass per fauna taxonomic relationships given in Marxsen et al. (2021). Prokaryote numbers were estimated by flow cytometry in samples stained with the fluorochrome Syto13 (Molecular Probes, Eugene, Oregon, U.S.A.) using the FACScalibur flow cytometer (Becton Dickinson, Franklin Lakes, New Jersey, U.S.A.) as detailed in Gasol & Giorgio (2000).

Multivariate analyses, i.e. nonmetric Multidimensional Scaling (NMDS), was calculated from the faunal assemblages. Since there were many zeros in the faunal assemblages’ data, the distance matrix for the NMDS was calculated using a zero-adjusted Bray–Curtis coefficient (Clarke et al., 2006), after Naupliae abundances were excluded. Since well assemblages with only this artificial abundance fell into one coordinate, they were drawn apart for visualization of hydrogeological provinces and were encircled with a grey circle. Environmental fitting of variables to the NMDS was done by the “envfit” procedure in the vegan package (Oksanen et al., 2022). In parallel to temperature, redox potential, CO_2_, HCO_3_^-^, and silica, further chemical and physical characteristics were chosen for the correlation with organism data. We focused on those that were measured most frequently, are most telling about hydrogeological conditions, and/ or are known to limit organisms: alkalinity, COD, Ca, Cl, DO, DOC, DRP, EC, F, Fe (dissolved), hardness, Mg, Mn, NH_4_^+^, NO_2_^-^, NO_3_^-^, Na^+^, SO_4_^2-^, pH. Also, their standard deviations, i.e. variability, and the slopes of the linear trends over time (see above) were included in the analyses, as well as abundances of prokaryotes and geographic coordinates. The variability that standard deviations indicates, may be due to interannual variability, or to the trend over time. If the standard deviation consists of both interannual variability and trend over time, it may lead to the system passing phase-tipping points (Alkhayuon et al., 2023). This meant 52 individual regressions. With that number of tests, the likelihood that some of the tests might by chance be significant (“false positive”), or by chance insignificant (“false negative”), increases, because of multiple testing. To compensate for such effect, the *p*-values were Benjamini-Hochberg-Yekutieli-corrected for multiple testing (Benjamini & Hochberg, 1995). Only envfit regressions with an adjusted *p*-value below 0.05 were marked as significant. All analyses, where not marked otherwise, were performed using R (R Core Team, 2023).

## Results

The averages, minima and maxima of those variables that are most relevant for weathering are listed in Table 1. In the lower part, the respective statistics are given separately for shallow, i.e. high exchange, and deep, i.e. low exchange, wells.

**Table 1:** Measured values for the most important variables, average, minima, and maxima for the total data set, and separately for deep, i.e. low exchange, and shallow, i.e. high exchange wells.

| Measured values |  | Temperature (°C) | pH | Redox (mV) | DO (mg / L) | pCO<sub>2</sub> (µatm) | HCO<sub>3</sub><sup>-</sup> (mg / L) | Si (mg / L) |
| --- | --- | --- | --- | --- | --- | --- | --- | --- |
| Total | Average | 10.25 | 6.42 | 36.43 | 2.12 | 46506 | 126.64 | 7.62 |
|  | Minimum | 2.70 | 4.40 | -181.00 | 0.01 | 0 | 0 | 0.03 |
|  | Maximum | 21.60 | 12.40 | 963.00 | 12.70 | 4641203 | 521.00 | 28.59 |
| Shallow | Average | 10.45 | 6.52 | 64.07 | 2.51 | 42516 | 165.20 | 6.17 |
|  | Minimum | 2.70 | 4.40 | -175.00 | 0.01 | 0 | 0 | 0.03 |
|  | Maximum | 21.60 | 8.40 | 436.00 | 12.30 | 4641203 | 521.00 | 28.59 |
| Deep | Average | 10.11 | 6.35 | 16.98 | 1.85 | 44974 | 98.72 | 8.67 |
|  | Minimum | 2.80 | 4.50 | -181.00 | 0.05 | 0 | 0 | 0.18 |
|  | Maximum | 16.80 | 12.40 | 963.00 | 12.70 | 800483 | 112.00 | 27.44 |

The developments over time of those variables most relevant for weathering are shown in SI 4, Figures S4 to S9, and Figure 1, and the full statistics are given statistics in Table S3. The slopes of the trends over time of those variable most influential in weathering are summarized in Table 2. In the lower part, the respective statistics separately for shallow and deep wells are also given. Only one well showed a slight insignificant decrease in Si values (shallow well VP1854), while values in all other wells increased significantly. In contrast, trends in the other variables varied, with increases and decreases regardless of whether they were deep or shallow.

**Table 2:** Slopes of the trend lines over time for the most important variables, average, minima, and maxima for the general trends and separate for deep, i.e. low exchange, and shallow, i.e. high exchange wells.

| Slope over time |  | Temperature (°C y<sup>-1</sup>) | pH (y<sup>-1</sup>) | Redox (mV y<sup>-1</sup>) | DO (mg L<sup>-1</sup> y<sup>-1</sup>) | pCO<sub>2</sub> (µatm y<sup>-1</sup>) | HCO<sub>3</sub><sup>-</sup> (mg L<sup>-1</sup> y<sup>-1</sup>) | Si (mg L<sup>-1</sup> y<sup>-1</sup>) |
| --- | --- | --- | --- | --- | --- | --- | --- | --- |
| Total | Average | 0.02 | 0 | -0.37 | -0.04 | 256.12 | 0.21 | 0.19 |
|  | Minimum | -0.29 | -0.03 | -6.74 | -0.42 | -2667.56 | -8.17 | -0.01 |
|  | Maximum | 0.17 | 0.02 | 10.25 | 0.12 | 4392.34 | 4.06 | 0.54 |
| Shallow | Average | 0.02 | 0 | 0.47 | -0.07 | 565.17 | 0.28 | 0.19 |
|  | Minimum | -0.29 | -0.03 | -4.70 | -0.42 | -1746.98 | -8.17 | -0.01 |
|  | Maximum | 0.12 | 0.02 | 10.25 | 0.10 | 4392.34 | 4.06 | 0.43 |
| Deep | Average | 0.02 | -0.01 | -1.00 | -0.02 | 35.38 | 0.16 | 0.19 |
|  | Minimum | -0.06 | -0.03 | -6.74 | -0.16 | -2667.56 | -1.76 | 0 |
|  | Maximum | 0.17 | 0.01 | 3.83 | 0.12 | 3759.66 | 2.69 | 0.54 |

**Figure 1:**
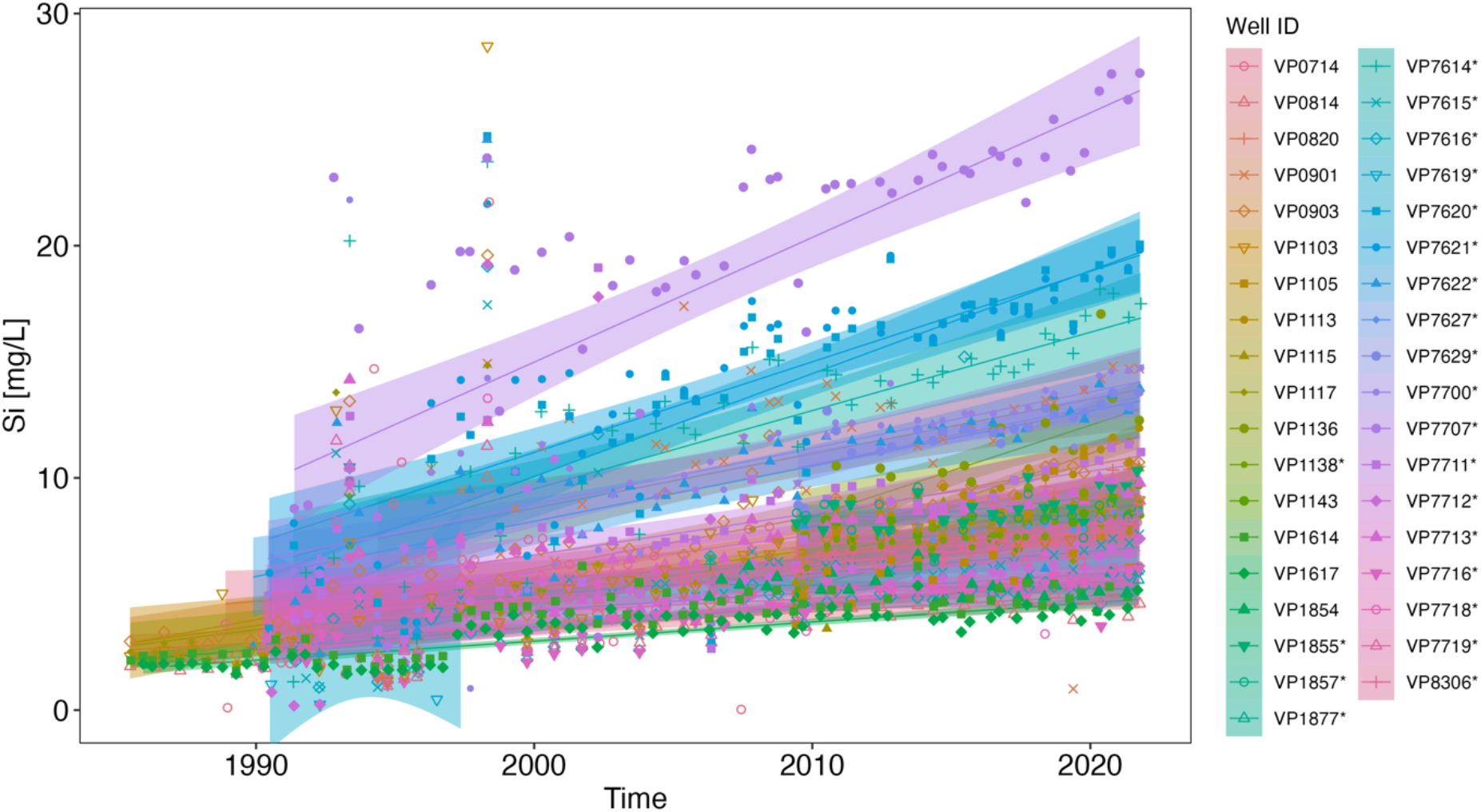
Silica, reported as Si concentration, over time in the wells that were sampled for fauna. Measurements, linear trend (in darker colour), and confidence interval (in lighter colour around the linear trend lines) are coloured according to the well. Well IDs of deep wells are marked with a star.

Molar ratios of Na/Ca versus Mg/Ca indicated that samples were closely related to value ranges known for silicate rock weathered water (Gaillardet et al., 1999; Négrel et al., 1993), as shown in Supplementary Information SI 6, Figure S10. While almost all wells can be considered to tap silicate rock type aquifers, the actual mean values of silica of the previous three years, and the average trend over the observation period, depended on the hydrogeological zone (SI 6, Fig. S10b). The redox potential and DO were lower in deep, than in shallow, quaternary wells (Wilcoxon rank sum test with continuity correction: W= 148521, p-value < 0.001, and W = 224719, p-value < 0.001, respectively).

Fauna was not found in all wells (SI 7, Fig. S11). There was no obvious geographical or hydrogeological pattern with which fauna was found – wells where high numbers were found were often close to wells where no fauna was found. Deep wells harboured fauna, and in contrast, not all shallow wells harboured fauna (compare the map in SI 7 and the results of the test in SI 8 with the map in SI 1, Fig. S1).

Fauna did not directly depend on prokaryote numbers (Figure 2), neither for abundances nor for biomass. Neither the regressions separated in shallow, nor those for deep aquifers, were significant (SI 9, Fig. S13).

**Figure 2:**
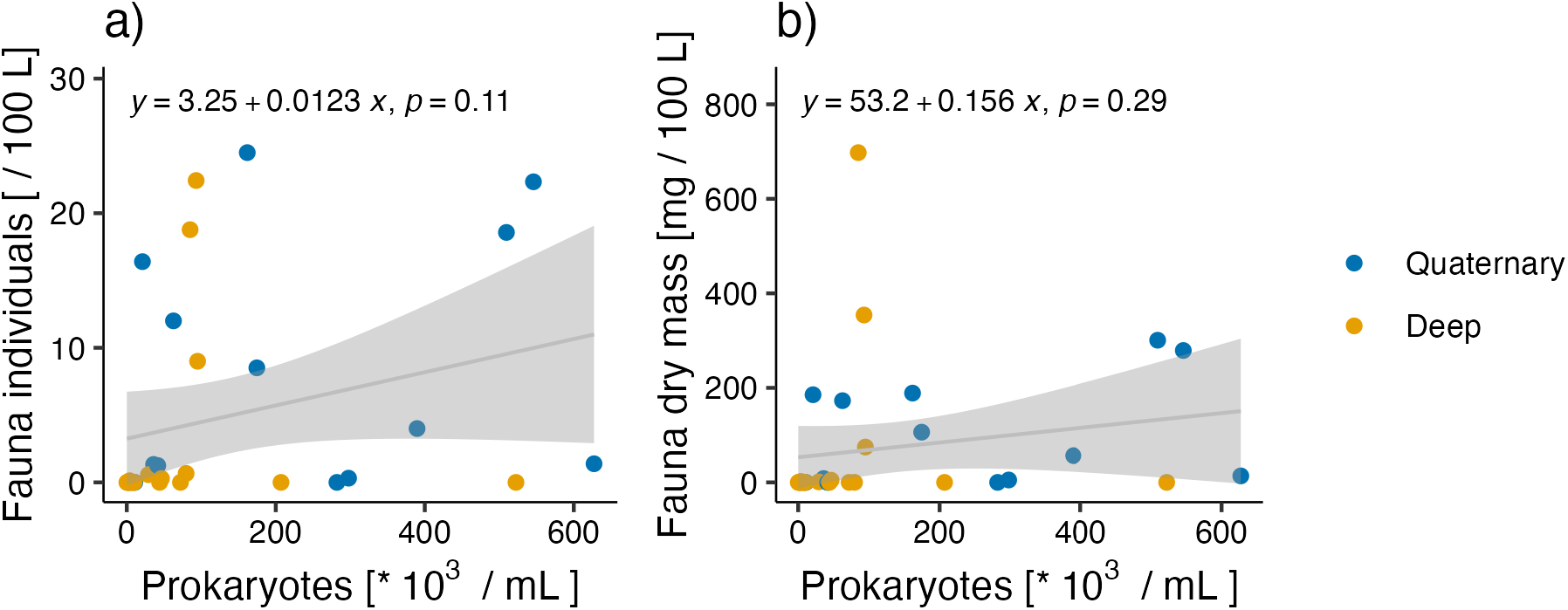
Fauna abundances (a) and fauna biomass (b) in dependence on prokaryote numbers. Given are the individual measurements (dots), the regression line (grey line), and the confidence interval (lighter grey area around the regression line), the regression equation (“y = … “), and the significance level p. For separate fits per aquifer type see Supplementary Information SI 8, Fig. S11.

Fauna individuals and biomass did not show a dependency on temperature, but highest abundances and biomasses seemed to be reached at temperatures between 10 and 11 °C in the shallow aquifer, and between 9 and 10 °C for the deeper aquifers (SI 9, Fig. S14 a-d). For prokaryotes, the temperature which yielded the highest cell numbers was in the low and high ranges for the deep aquifers, and close to 11 °C for prokaryotes from the shallow aquifers (SI 9, Fig. S14 e, f).

The current redox potential was positively correlated with abundances and biomass for all aquifers taken together (SI 9, Figure S15 a, c), and was highly positively related for deep, but not for shallow aquifers (SI 9, Figure S15 b, d). The prokaryote numbers were not related to redox potential, neither for all aquifers taken together, nor separately for deep and shallow aquifers (SI 9, Figure S15 e, f). The patterns for the current dissolved oxygen (DO) concentrations were similar to those for redox potential (SI 9, Figure S16), although redox potential and DO were only slightly correlated (Spearman rank correlation r_s_ = 0.51). Neither CO_2_ nor HCO_3_^-^ concentrations and organisms were correlated (SI 9, Figure S17, S18).

As hypothesized, fauna and prokaryotes reacted in a similar way to increasing silica concentrations (Figure 3). In contrast to the expectation, fauna abundances (Figure 3a, b) and biomass (SI 9, Figure S19), as well as prokaryote numbers (Figure 3c, d) were negatively correlated to silica concentrations. For fauna individuals, this was due to the values from shallow, quaternary wells (Figure 3b).

**Figure 3:**
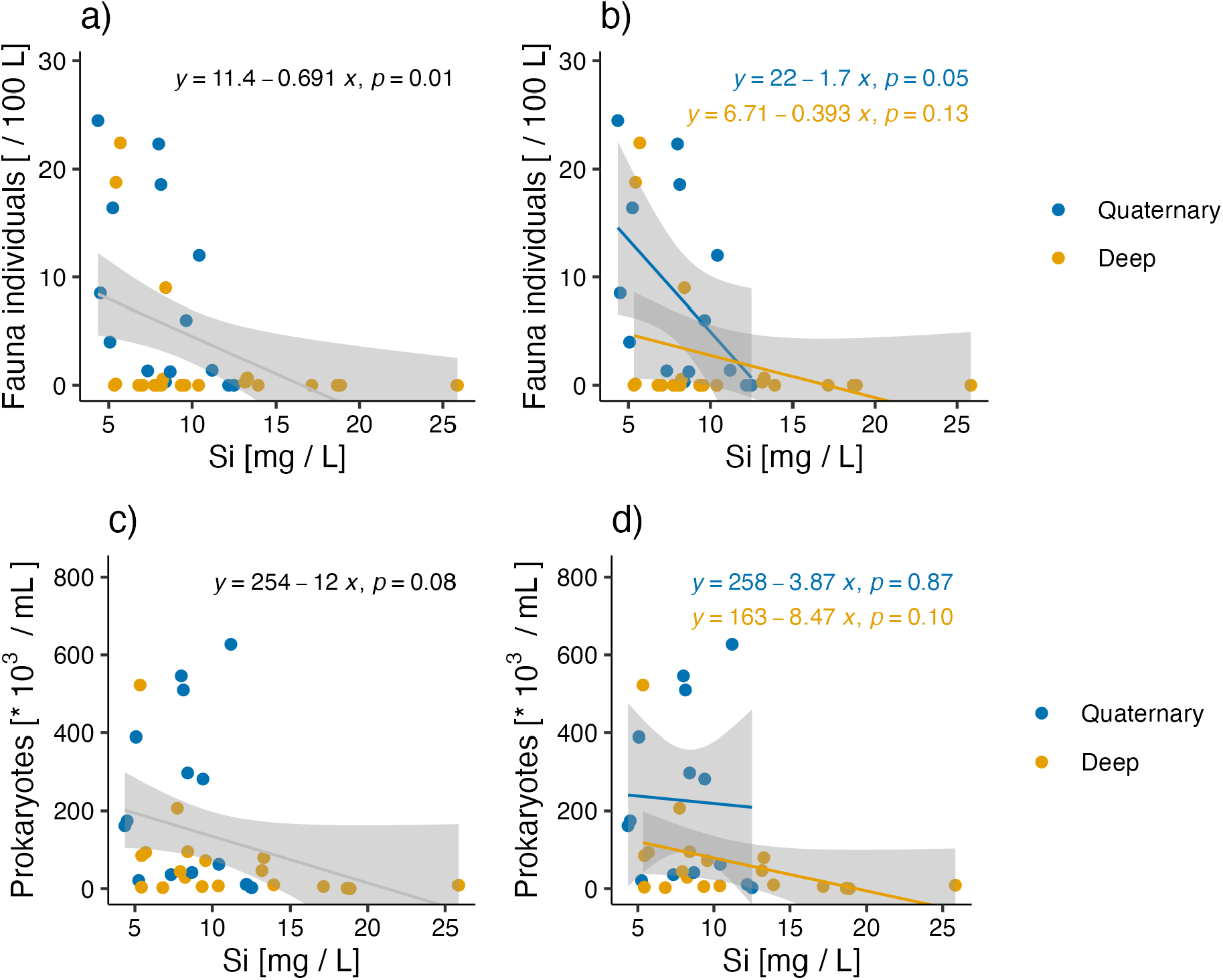
Organisms in dependence on silica concentrations. Top row: fauna abundances. a) Total trends; b) trends separately for deep (yellow) and shallow (blue) aquifers; Bottom row: prokaryotes. c) Total trends; d) trends separately for deep (yellow) and shallow (blue) aquifers. Plots for fauna biomass are in the SI 9, Figure S19. The equations for the trends are given above the respective plot.

Like the current measured values for temperature, and CO_2_ and HCO_3_^-^ concentrations, also the trends of temperature (SI 9, Figure S20), redox potential (SI 9, Figure S21), DO (SI 9, Figure S22), CO_2_ (SI 9, Figure S23), and HCO_3_^-^ (SI 9, Figure S24) concentrations were not related to organisms biomass and numbers, neither for fauna nor for prokaryotes. Fauna individuals and biomass slightly, but insignificantly, decreased with increasing slopes in silica change over time (Figure 4a, b; SI 9, Figure S25 for fauna biomass). Prokaryote numbers, in contrast, in total slightly increased (Figure 4c), but significantly decreased in deep wells (Figure 4d).

**Figure 4:**
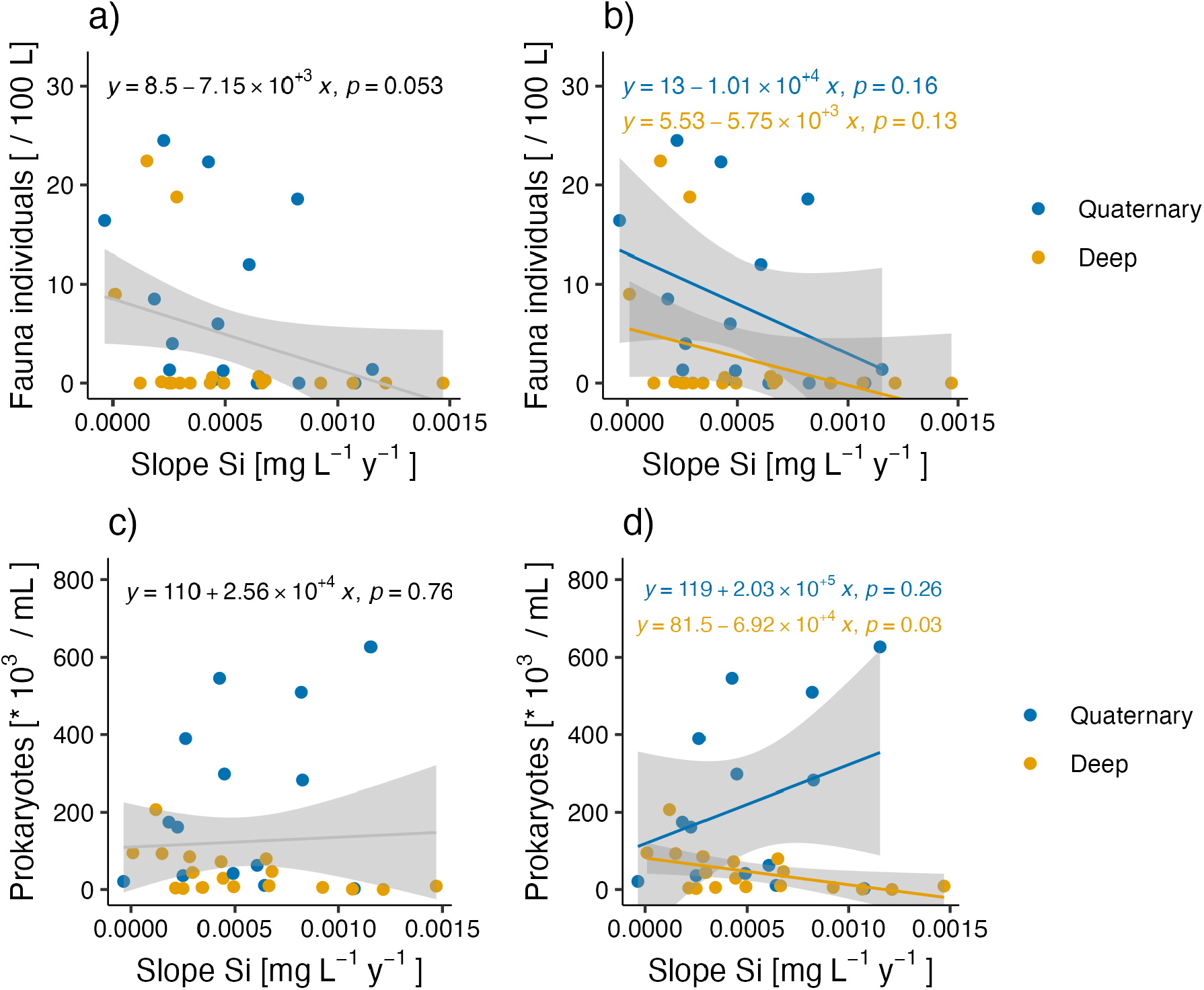
Dependence of fauna individuals (a, b), and prokaryotes (c, d) on the slope in silica concentration change over time. Regressions are drawn separately for quaternary and deep aquifers in b) and d). Plots for fauna biomass are in the SI 9, Figure S25.

Nonmetric Multidimensional Scaling (NMDS) showed that there was no pattern in fauna favouring certain hydrogeological provinces over others, or fauna assemblages being fundamentally different in some hydrogeological provinces – the colours for the hydrogeological provinces are well distributed within Figure 5a, and wells where no fauna was found were situated in diverse hydrogeological zones. These wells where no fauna was found were characterized by high Si and dissolved iron (“Fe_d”) concentrations, and were negatively correlated with Redox potential (“Eh”), nitrate (“NO3_N”; Figure 5b, SI 10, Table S4), and the variability in nitrate (“sd_NO3”; Figure 5b, SI 10, Table S4). Independently of these concentrations, the variability in temperature (“sd_T”) and dissolved oxygen concentrations (“DO”) shaped faunal assemblages, particularly in the Quaternary of the Otava River, the Sokolov Basin, Budweis Basin, and Crystalline Upper Vltava (Figure 5b).

**Figure 5:**
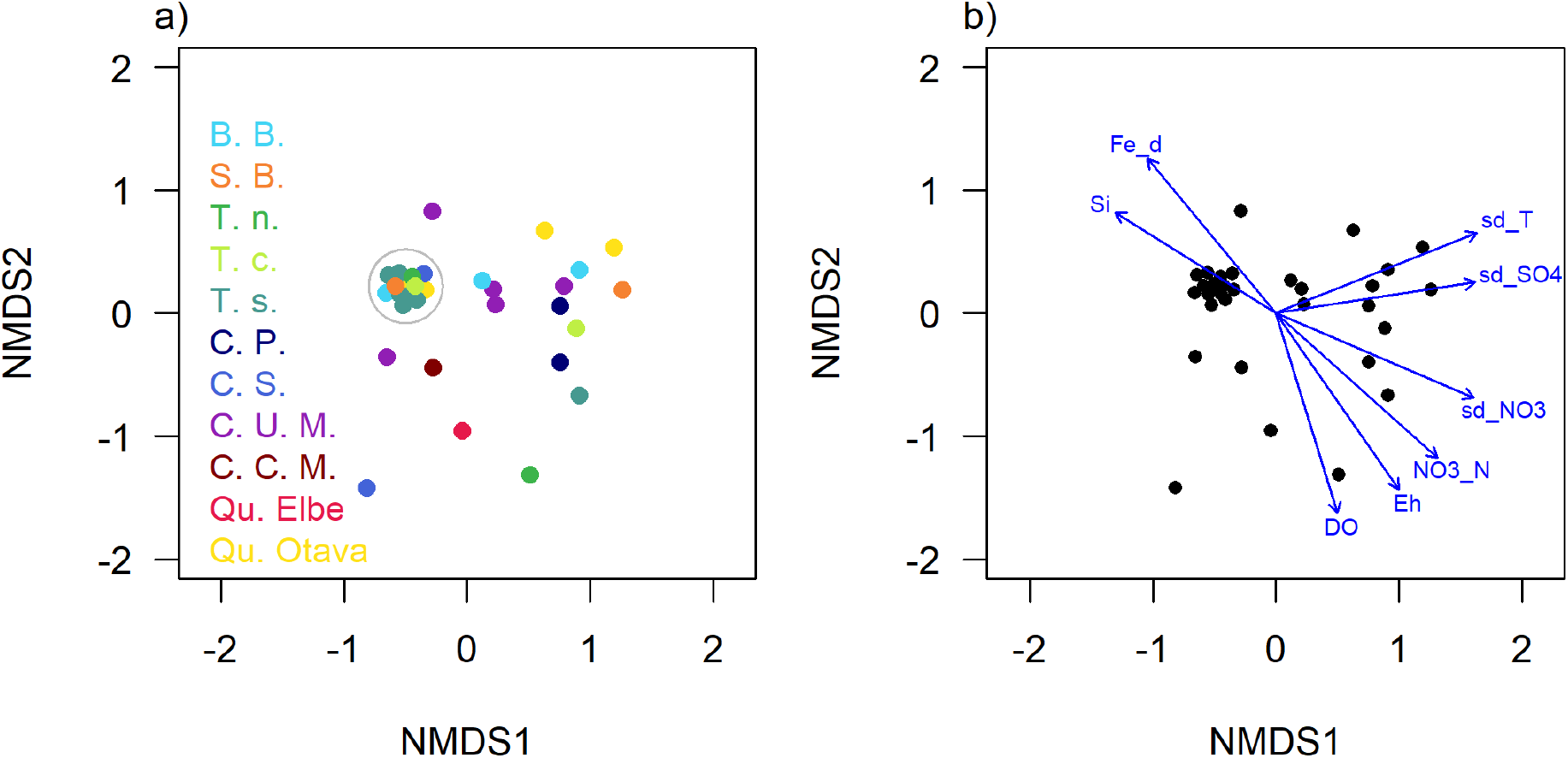
Nonmetric Multidimensional Scaling (NMDS) of the zero-adjusted faunal assemblage. Since well assemblages with only this artificial abundance fell into one coordinate, they were drawn apart for visualization of hydrogeological basins and were encircled with a grey circle. a) Sites coloured according to hydrogeological province. “B. B.” = Budweis basin, “S. B.” = Sokolov basin, “T. n.” = Třebon basin north, “T. c.” = Třebon basin centre, “T. s.” = Třebon basin south, “C. P.” Crystalline Proterozoic, “C. S.” = Crystalline Smrčín and southern parts of Ore Mountains, “C. U. M.” = Crystalline Upper Vltava, “C. C. M.” = Crystalline Central Vltava, “Qu. Elbe” = Quaternary of the Labe to the Vltava, “Qu. Otava” = Quaternary River Otava. b) The same plot as in a), but overlain by significant correlations with chemical and physical properties, identified by the envfit procedure. “sd_” = standard deviation of the variable, “T” = temperature; “NO2_N” = nitrite nitrogen, “SO4” = sulphate, “NO3” or “NO3_N” = nitrate nitrogen, “COD” = chemical oxygen demand, “proks” = prokaryotes, “Si” = silica, “NH4_N” = ammonia nitrogen, “Fe_d” = dissolved iron; “_slope” = slope over time as yearly slope, “DO” = dissolved oxygen. Labels in b) slightly pulled apart to increase readability.

## Discussion

Temperature and CO_2_ levels may be seen as the main drivers of a changing climate in aqueous systems. However, in the investigated aquifers neither did they show clear trends, nor did they correlate with organisms. The clearest trend was exhibited by silica, i.e. the result of weathering which was likely induced by increased temperature and CO_2_. Both silica and redox potential showed the clearest correlation with organisms. These are likely indirect effects, as will be clarified throughout this discussion.

### General physical and chemical conditions in the study area

The temperatures, pH, redox potential, and concentrations of DO, CO_2_, HCO_3_^-^ and silica were within the ranges typical for groundwater, with some values of HCO_3_^-^ being at the upper end of usual values (Fitts, 2012). As expected, the redox potential was lower in deeper, low-exchange aquifers which may have undergone various cycles of biogeochemical reactions, reducing oxidized compounds. This underlines the importance of studying deep and shallow aquifers separately. The high HCO_3_^-^ values could be due to increased CO_2_ recharge, as observed by Macpherson et al. (2008). The estimated CO_2_ itself was up to three orders of magnitudes higher in this study than the values given in Macpherson et al. (2008). The pH which was lower in the Czech aquifers than in the limestone aquifer studied by Macpherson et al. (2008) would have moved the carbonate equilibrium towards higher CO_2_ at the expense of HCO_3_^-^ in most aquifers.

### Trends in physical and chemical conditions

In contrast to the first hypothesis, and despite clear increases in air temperature over the past 40 years in the Czech Republic (Zahradníček et al., 2021), groundwater temperature only increased in ca. 80% of aquifers over the previous ca. 30 years. However, groundwater temperature is the result of conduction and advection. Heat is conducted along soil and aquifer material along the temperature gradient according to the thermal conductivity of the material in question. The transport of heat with water recharging from the surface into groundwater is called heat advection. When trending over time, one has to take into account the depth of sampling, seasonality (i.e. more than two samples per year in order to increase the chance to see the minima and maxima; Benz et al., 2017), and the properties of the geological material (Davies, 2013). The idea that groundwater temperature just follows surface temperature, is too simplistic – several processes are involved (Rau et al., 2015). In neighbouring Bavaria, Hemmerle & Bayer (2020) also found one out of 32 groundwater wells to have decreased in temperature between 1994 and 2019, although air temperature had increased considerably.

Likewise, CO_2_ and HCO_3_^-^ concentrations did not increase in all aquifers in contrast to the hypothesis – some aquifers showed decreasing trends. Groundwater is often supersaturated in CO_2_, particularly where it is under pressure (Bailey, 1991), or where there was considerable microbial production (Chapelle et al., 1987), and thus, import beyond saturation might not be physically possible. However, rivers are also often supersaturated and still showed an increasing trend in the recent decades (e.g. Raymond & Cole, 2003), attributed to increased atmospheric CO_2_, increased water flow, and changing land use in the Mississippi River. The reason for the lack in clear patterns in Western Czechia might lie in the fact that HCO_3_^-^ is in a pH-dependent equilibrium with CO_2_ and CO_3_^2-^ and the HCO_3_^-^ concentration may change in response to changes in CO_2_ import into groundwater, pH-dependent transformation, carbonate precipitation, and uptake by organisms. Kump et al. (2000) assumed that soil pCO_2_ was determined by microbial activity, and not directly by the atmospheric concentrations. This may hold true for some groundwater aquifers where microbial productivity is high, as well.

However, Kump et al. (2000) also concluded indirect effects of atmospheric pCO_2_ on the soil, via the plant productivity which increases with pCO_2_. This increased plant productivity may have effects on at least the shallow groundwater. Also, since local heterogeneity is probably much higher than usually accounted for (Schmidt et al., 2017), it is not unreasonable to expect that locally, on the micro scale, a groundwater pore becomes undersaturated in CO_2_ due to e.g. intense small-scale chemolithoautotrophy using CO_2_ as a reaction partner. On this small scale, CO_2_ might then be transferred from the surface into groundwater. The overall supersaturation might thus increase on average, and this might have happened in some aquifers where CO_2_ levels did overall increase. Decreasing CO_2_ concentrations in other aquifers, while atmospheric CO_2_ increased, probably involved complex chemical reactions. These processes of import, transformation, or export were not measured themselves during the monitoring presented here – this would have required flux meters, isotopes, and related techniques.

In contrast to CO_2_ and HCO_3_^-^ concentrations, silica concentrations rose in all but one aquifer, and the rise was significant (Figure 1, Supporting Information Table S2). This is most likely a consequence of silicate weathering which would have transformed CO_2_ to HCO_3_^-^ (Walker et al., 1981; Zhang et al., 2021), another possible explanation for the lack in patterns in CO_2_ and HCO_3_^-^ concentrations (for a more complete discussion, see Supplementary Information 1). It is thus possible that the CO_2_ import had temporarily increased in more aquifers than where it is now measurable, but that the CO_2_ was consumed in weathering, and we only look at the net CO_2_ concentrations, but can estimate the extent of increased weathering by the increase in silicate. Macpherson et al. (2008) observed similar patterns of CO_2_ partly being consumed by weathering in a limestone Konza prairie aquifer.

In agreement with the second hypothesis, shallow and deep aquifers differed in the direction and steepness of trends, particularly trends of silica concentrations, with deep aquifers harbouring the one insignificant decrease observed, but generally a steeper increase than in shallow aquifers (Supplementary Information Table S2). The data series only spans up to 40 years, and many wells in this investigation had only been monitored for about a decade. It is possible that there had been an increase in silicate weathering in shallow aquifers until one or two decades ago and – like the effects from acidification – this increase now equilibrates to a new state. One indicator for such a pattern is that the current silica concentrations are higher in shallower aquifers than in deep ones, despite the presumably shorter residence and younger age of the former. Usually, weathering rates depend inherently on the residence time (Bakalowicz, 1994; Maher, 2010) and silica concentrations are known to reflect the age of the aquifer (Marçais et al., 2018). Shallow aquifers with a lower age but higher silica concentrations may reflect an interim stage that the deeper groundwater will pass beyond, given time. There is no telling what the new equilibrium for deeper aquifers will be, and what this will mean for organisms.

While increases in CO_2_ and HCO_3_^-^ are known from rivers (Macpherson & Sullivan, 2019), increases in silica are rarely mentioned and net increases might not have occurred in surface waters (e.g. Macpherson & Sullivan, 2019). This is due to the fact that silicate is used by stream organisms, mainly diatoms (Lampert & Sommer, 2007), which do not grow in groundwater. Rivers where such groups as diatoms grow are thus not ideal for studying the effect of weathering on silica concentrations since the newly dissolved silica may be used up immediately. It makes more sense to study weathering effects directly in groundwater.

### Did organisms reflect the current physical and chemical conditions?

In contrast to the third hypothesis, neither aquifer temperature, nor recent CO_2_ and HCO_3_^-^ concentrations, were significantly correlated with organism numbers, neither for fauna nor for prokaryotes (Fig. 3; SI 8, Fig. S13 S16). Thus, temperature, CO_2_ or HCO_3_^-^ neither seemed to harm nor promote organisms. Regarding temperature, this apparently contrasts the findings by Brielmann et al. (2009) and Spengler & Hahn (2018), but in the cited references, wells were compared that were under varyingly intense anthropogenic influence, while the wells sampled in this study, were not directly influenced by human activity beyond global influences such as rising CO_2_ and increasing land use – wells were neither in highly industrialized areas, nor in high-productivity agricultural areas or large cities. This might mean that the increases of air temperature have not begun to impact on groundwater ecosystems yet, or these effects were cancelled out by other factors and developments.

The positive relationship between redox potential and DO on the one hand, and fauna on the other hand, were to be expected since most multicellular fauna require at least minute concentrations of dissolved oxygen (Malard & Hervant, 1999). In addition, if the redox potential, which was not strongly correlated to DO, indeed was an indicator for the vulnerability of the aquifers or for the degree of exchange with the surface (Meckenstock et al., 2015), then the positive relationship between redox potential and fauna indicates the dependence of fauna on exchange, as postulated earlier (Schmidt & Hahn, 2012). Contrary to expectation, prokaryote abundances were neither positively correlated with DO nor redox potential. Microbial productivity is usually directly positively related to redox conditions, since more oxidized compounds provide more energy in biogeochemical reactions. The reason for the apparent lack of the expected positive relationship between redox potential and microorganisms may be due to the measured microorganism numbers not being representative for the actual microbial production. The expected higher biomass might have been grazed by fauna to a higher degree. Fauna itself then increased in numbers and biomass along the redox gradient. Fauna may have thereby reduced prokaryote stock to a minimum below which prokaryote production would not support fauna production. We propose that fauna relied on prokaryote growth more in deep wells than in shallow wells where exchange with the surface might deliver sufficient palatable food sources (but see the discussion in Karwautz et al., 2022, on alpine, low organic input aquifers). This may indicate that deeper wells are generally more high-quality carbon and energy-limited than the shallower wells where exchange with the surface is bound to be more intense. We suggest that chemolithoautotrophic carbon fixation, which is likely to have been important in the deep, low-exchange wells, is indeed an important carbon and energy source for food chains as coined by Overholt et al. (2022), but all the more so, the lower the exchange with the surface, as shown in Herrmann et al. (2017, 2020). The observed prokaryote numbers may thus represent a base production below which ecosystem productivity does not go, unless disturbed e.g. by agriculture (Marxsen et al., 2021).

The lack in correlation between organisms and CO_2_ is surprising, given that the values ranged from 0 to 4 864 077 µatm, which is almost 60 times as high as the concentration tested by Ramaekers et al. (2022) who found lethal effects for daphnids, ostracods, and rotifers at 83 000 µatm CO_2_. Already at 18 000 µatm, Weiss et al. (2018) found that the increased CO_2_ impaired the ability to sense predators in surface zooplanktonic *Daphnia* and their ability to form the inducible defences in the form of helmets. Increased CO_2_ thus rendered the daphnids more prone to predation. The authors predicted impacts on the whole food chain. Crustacea dominate groundwater fauna biomass and it is thus not unreasonable to expect similar impairments to groundwater fauna where CO_2_ concentrations increase more clearly. However, groundwater taxa are characterized by different traits than surface water organisms (Hose et al., 2022), and it is thus possible that groundwater organisms are better adapted to higher CO_2_ concentrations than surface water organisms. The lack in correlations between temperature and inorganic carbon concentrations (CO_2_ and HCO_3_^-^) on the one hand and organisms on the other hand might point to the single effects on organisms being cancelled out by each other. Tracer and physiological experiments might shed light into the interactions between temperature and inorganic carbon on organisms.

The negative effect of silica on organisms was unexpected and unlikely to be a direct effect, since silica is not known to be harmful to organisms – on the contrary, it is a micronutrient for e.g. diatoms, foraminifera, and radiolaria (Lampert & Sommer, 2007; Moore et al., 2013) – which, however, do not occur in groundwater. One explanation for the negative trend between fauna abundances and silica may be that silica reflects the age of the aquifer (Marçais et al., 2018). Fauna may thus be negatively correlated with the age of the aquifer. The reason for this may lie in the quality of carbon, in terms of degradability, decreasing over time, which is not reflected in the concentrations of DOC or TOC (Hofmann et al., 2020). However, the patterns of silica having risen to higher levels in shallow aquifers and not rising as much anymore in shallow aquifers than in deep aquifers, points to another possible explanation, i.e. weathering. The increasing silica concentrations indicate the increased weathering. Weathering itself is not harmful to fauna – some taxa are actively involved in silica weathering without any obvious sign of damage (Vicca et al., 2022) or even rely on it as a fertilizing process (Kelland et al., 2020). In fact, Schwartzman & Volk (1989) estimated that without biogenic weathering, the earth would be tens of degrees warmer, than it is. However, through weathering, other chemical compounds might have become bioavailable (Rue & McKnight, 2021) that might have been harmful to organisms (Jones & Bennett, 2014; Vicca et al., 2022), such as aluminum which is a common companion in silica minerals (Amann et al., 2020). Aluminum is toxic to many organisms in the microgram per Litre range (Botté et al., 2022).

Liu et al. (2016) proposed that the higher content of silica in diatoms led to them being less nutritional, harder to digest, leading to denser fecal pellets and lower fecundity rate in crustacea. Thus, a further explanation may lie in fauna digesting the weathered sand grains which are enriched in silica following the weathering. The results by Liu et al. (2016) might imply that silica wreaks havoc in the digestion tracts of crustacea. However, the proximate factor may not be silica but the cell wall thickness and rigidity. Liu et al. (2016) compared fauna’s productivity when grown on two different populations of a diatom species, grown at high and low light intensity, with the low light intensity population harbouring higher silica in the cell walls. However, the cell wall may have differed also in e.g. poly-unsaturated fatty acids (PUFA) which are decisive for many body functions of the grazer, including reproduction (Brett et al., 2017). Temperature and CO_2_ have been found to influence the fatty (FA) and amino acid (AA) composition in primary producers (Bermúdez et al., 2015). E.g. high CO_2_-grown diatoms had less PUFA than controls (Bermúdez et al., 2015). Thus, high silica concentrations may be the indicator for a high degree of CO_2_ having recharged into the water which may have led to decreased food quality, rather than silica itself being harmful to fauna.

### Do organisms reflect the trends in physical and chemical conditions?

Largely, the trends of temperature, CO_2_ and HCO_3_^-^ over time were not reflected in nowadays organism numbers in contrast to hypothesis 4, nor in the distribution of species in among the aquifers, in contrast to hypothesis 5. However, neither do we know when exactly the changes in groundwater began, i.e. how much change had already taken place before the monitoring started, nor what the populations were before the changes began. Also, we do not know how much more the aquifers will change and what that will mean to organisms, i.e. whether the ecosystems are at risk. It was hypothesized that fauna would be negatively correlated to an increase in silica concentrations, i.e. anthropogenically accelerated silica outflow rate, because that will reflect change which the groundwater organisms might not be able to withstand. However, this correlation was not significant, in contrast to the correlation with the current concentrations (see previous paragraphs).

One reason for CO_2_ neither showing temporal trends, nor being clearly related to organisms, may be that the additional CO_2_ import into groundwater is divided into pathways – parts of it may have been involved in weathering of silicate rock, while another part may have (and maybe increasingly so over the past decades) taken part in chemolithoautotrophic processes.

### The geological thermostat

The uptake of CO_2_ during bio-erosion of geological structures, which leads to increased aqueous concentrations of silica, reduces the greenhouse gas CO_2_ in the atmosphere and thus reduces global air temperatures. This phenomenon has been called the geological thermostat (Li et al., 2016). On the long-term time scales, but also on the shorter-term, this extends habitability of the planet Earth (Brantley et al., 2023). However, if one of the side effects of this thermostat is the loss of organisms and loss of the ecosystem functions they perform, this may negatively impact on habitability of the planet. We do rely on self-cleaning within aquifers and rivers for drinking water production (Griebler & Avramov, 2015). E.g. Prague, which receives most of the water studied in this contribution, is among the cities that are groundwater-dependent for their drinking water; in the case of Prague, drinking water production relies to 25% on groundwater (Vejvodová et al., 2006).

## Conclusion

Climate change in terms of increased temperatures was not the main driver for organism occurrence and fauna communities. However, the impacts from climate change are still in progress and patterns might change when surface temperatures have increasingly reached subsurface realms. More important than the recent temperature, seems to have been the effect of increased CO_2_ the net concentrations of which did not increase measurably in the studied wells, but which seems to have been responsible for increased silicate weathering in the aquifers the concentrations and increases of which correlated negatively with organisms. That the process is on-going, can be deduced from the observation that the silica concentrations were partly higher in shallow aquifers with presumably a younger age, while one might have expected that silica is highest in oldest, i.e. deepest aquifers. This seems to indicate that the aquifers are still in development towards a new equilibrium. It remains to be seen what the new equilibrium for deeper aquifers will be, and what this will mean for organisms and ecosystem services that we rely on for e.g. drinking water production. It is also likely that while temperature and CO_2_ on their own did not correlate with organism abundances, their interaction might have done. Lastly, this study focused on standing stocks. Transfer rates of the abiotic and biotic components are necessary to further clarify processes.

## Acknowledgements

SIS acknowledges funding through MEMOBIC [EU Operational Programme Research, Development and Education No. CZ.02.2.69/0.0/0.0/16_027/0008357], by The Ministry of Education, Youth and Sports of the Czech Republic [grant number CZ.02.1.01/0.0/0.0/16 025/0007417], and from the TAČR KAPPA project No. 2020TO01000202 funded by the Norway Grants. We are grateful to many colleagues who participated in this research, particularly the staff of ALS who were very helpful in enabling sampling for organisms, to Mark Cuthbert for discussions on temperature trends in groundwater, to Robert Lehmann for discussions on weathering, to Jiři Kopáček for discussions on silica, and to Martina Herrmann for discussions on chemolithoautotrophy.

## References

Alkhayuon, H., Marley, J., Wieczorek, S., & Tyson, R. C. (2023). Stochastic resonance in climate reddening increases the risk of cyclic ecosystem extinction via phase-tipping. Global Change Biology, 29(12), 3347–3363. 10.1111/gcb.16679

Amann, T., Hartmann, J., Struyf, E., de Oliveira Garcia, W., Fischer, E. K., Janssens, I., Meire, P., & Schoelynck, J. (2020). Enhanced Weathering and related element fluxes – a cropland mesocosm approach. Biogeosciences, 17(1), 103–119. 10.5194/bg-17-103-2020

Aquilina, L., Stumpp, C., Tonina, D., & Buffington, J. M. (2023). Chapter 1— Hydrodynamics and geomorphology of groundwater environments. In F. Malard, C. Griebler, & S. Rétaux (Eds.), Groundwater Ecology and Evolution (Second Edition) (pp.3–37). Academic Press. 10.1016/B978-0-12-819119-4.00014-7

Bailey, R. A. (1991). Chemistry of the Environment.

Bakalowicz, M. (1994). 4 Water geochemistry: Water quality and dynamics. In J. Gibert, D. L. Danielopol, & J. A. Stanford (Eds.), Groundwater Ecology (pp. 97–127). 10.1016/B978-0-08-050762-0.50011-5

Benjamini, Y., & Hochberg, Y. (1995). Controlling the False Discovery Rate: A Practical and Powerful Approach to Multiple Testing. Journal of the Royal Statistical Society: Series B (Methodological), 57(1), 289–300. 10.1111/j.2517-6161.1995.tb02031.x

Benz, S. A., Bayer, P., & Blum, P. (2017). Global patterns of shallow groundwater temperatures. Environ. Res. Lett., 12, 034005.

Bermúdez, R., Feng, Y., Roleda, M. Y., Tatters, A. O., Hutchins, D. A., Larsen, T., Boyd, P. W., Hurd, C. L., Riebesell, U., & Winder, M. (2015). Long-term conditioning to elevated pCO2 and warming influences the fatty and amino acid composition of the diatom Cylindrotheca fusiformis. PLOS One, 10(5), e0123945. 10.1371/journal.pone.0123945

Boerner, A. R., & Gates, J. B. (2015). Temporal dynamics of groundwater-dissolved inorganic carbon beneath a drought-affected braided stream: Platte River case study. Journal of Geophysical Research: Biogeosciences, 120(5), 924–937. 10.1002/2015JG002950

Botté, A., Zaidi, M., Guery, J., Fichet, D., & Leignel, V. (2022). Aluminium in aquatic environments: Abundance and ecotoxicological impacts. Aquatic Ecology, 56(3), 751–773. 10.1007/s10452-021-09936-4

Bou, C. (1974). Recherche sur les eaux souterraines. Les méthodes de récolte dans les eaux souterraines interstitielles. Ann. Spéolo., 29(4), 611–619.

Brantley, S. L., Shaughnessy, A., Lebedeva, M. I., & Balashov, V. N. (2023). How temperature-dependent silicate weathering acts as Earth’s geological thermostat. Science, 379(6630), 382–389. 10.1126/science.add2922

Brett, M. T., Bunn, S. E., Chandra, S., Galloway, A. W. E., Guo, F., Kainz, M. J., Kankaala, P., Lau, D. C. P., Moulton, T. P., Power, M. E., Rasmussen, J. B., Taipale, S. J., Thorp, J. H., & Wehr, J. D. (2017). How important are terrestrial organic carbon inputs for secondary production in freshwater ecosystems? Freshwater Biology, 1–21. 10.1111/fwb.12909

Brielmann, H., Griebler, C., Schmidt, S. I., Michel, R., & Lueders, T. (2009). Effects of thermal energy discharge on shallow groundwater ecosystems. FEMS Microbiology Ecology, 68(3), 273–286. 10.1111/j.1574-6941.2009.00674.x

Calmels, D., Galy, A., Hovius, N., Bickle, M., West, A. J., Chen, M.-C., & Chapman, H. (2011). Contribution of deep groundwater to the weathering budget in a rapidly eroding mountain belt, Taiwan. Earth and Planetary Science Letters, 303(1), 48–58. 10.1016/j.epsl.2010.12.032

Chapelle, F. H., Zelibor, J. L., Grimes, D. J., & Knobel, L. L. (1987). Bacteria in deep coastal plain sediments of Maryland: A possible source of CO2 to groundwater. Water Resources Research, 23(8), 1625–1632.

Clarke, K. R., Somerfield, P. J., & Chapman, M. G. (2006). On resemblance measures for ecological studies, including taxonomic dissimilarities and a zero-adjusted Bray-Curtis coefficient of denuded assemblages. Journal of Experimental Marine Biology and Ecology, 330, 55–80.

Cole, J. (2013). Freshwater in flux. Nature Geoscience, 6(1), Article 1. 10.1038/ngeo1696

Conant, R. T., Ryan, M. G., Ågren, G. I., Birge, H. E., Davidson, E. A., Eliasson, P. E., Evans, S. E., Frey, S. D., Giardina, C. P., Hopkins, F. M., Hyvönen, R., Kirschbaum, M. U. F., Lavallee, J. M., Leifeld, J., Parton, W. J., Megan Steinweg, J., Wallenstein, M. D., Martin Wetterstedt, J. Å., & Bradford, M. A. (2011). Temperature and soil organic matter decomposition rates – synthesis of current knowledge and a way forward. Global Change Biology, 17(11), 3392–3404. 10.1111/j.1365-2486.2011.02496.x

Danovaro, R., Dell’Anno, A., Pusceddu, A., Gambi, C., Heiner, I., & Kristensen, R. M. (2010). The first metazoa living in permanently anoxic conditions. BMC Biology, 8(1), n30. 10.1186/1741-7007-8-30

Davies, J. H. (2013). Global map of solid Earth surface heat flow. Geochemistry, Geophysics, Geosystems, 14(10), 4608–4622. 10.1002/ggge.20271

Fenchel, T., & Finlay, B. (2008). Oxygen and the spatial structure of microbial communities. Biological Reviews of the Cambridge Philosophical Society,83(4), 553–569. 10.1111/j.1469-185X.2008.00054.x

Fitts, C. R. (2012). Groundwater Science (Second Edi). Elsevier.

Friedlingstein, P., O’Sullivan, M., Jones, M. W., Andrew, R. M., Gregor, L., Hauck, J., Le Quéré, C., Luijkx, I. T., Olsen, A., Peters, G. P., Peters, W., Pongratz, J., Schwingshackl, C., Sitch, S., Canadell, J. G., Ciais, P., Jackson, R. B., Alin, S. R., Alkama, R., … Zheng, B. (2022). Global Carbon Budget 2022. Earth System Science Data, 14(11), 4811–4900. 10.5194/essd-14-4811-2022

Gaillardet, J., Dupré, B., Louvat, P., & Allègre, C. J. (1999). Global silicate weathering and CO2 consumption rates deduced from the chemistry of large rivers. Chemical Geology, 159(1), 3–30. 10.1016/S0009-2541(99)00031-5

Gasol, J. M., & Giorgio, P. A. del. (2000). Using flow cytometry for counting natural planktonic bacteria and understanding the structure of planktonic bacterial communities. Scientia Marina, 64(2), Article 2. 10.3989/scimar.2000.64n2197

Griebler, C., & Avramov, M. (2015). Groundwater ecosystem services: A review. Freshwater Science, 34(1), 355–367. 10.1086/679903

Hasler, C. T., Hannan, K. D., Jeffrey, J. D., & Suski, C. D. (2017). Valve movement of three species of North American freshwater mussels exposed to elevated carbon dioxide. Environmental Science and Pollution Research, 24(18), 15567–15575. 10.1007/s11356-017-9160-9

Hemmerle, H., & Bayer, P. (2020). Climate Change Yields Groundwater Warming in Bavaria, Germany. Frontiers in Earth Science, 8. 10.3389/feart.2020.575894

Herrmann, M., Geesink, P., Yan, L., Lehmann, R., Totsche, K. U., & Küsel, K. (2020). Complex food webs coincide with high genetic potential for chemolithoautotrophy in fractured bedrock groundwater. Water Research, 170, 115306. 10.1016/j.watres.2019.115306

Herrmann, M., Opitz, S., Harzer, R., Totsche, K. U., & Küsel, K. (2017). Attached and suspended denitrifier communities in pristine limestone aquifers harbor high fractions of potential autotrophs oxidizing reduced iron and sulfur compounds. Microbial Ecology,74(2), 264–277. 10.1007/s00248-017-0950-x

Herrmann, M., Rusznyák, A., Akob, D. M., Schulze, I., Opitz, S., Totsche, K. U., & Küsel, K. (2015). Large fractions of CO2-fixing microorganisms in pristine limestone aquifers appear to be involved in the oxidation of reduced sulfur and nitrogen compounds. Applied and Environmental Microbiology, 81(7), 2384–2394. 10.1128/AEM.03269-14

Hofmann, R., Uhl, J., Hertkorn, N., & Griebler, C. (2020). Linkage between dissolved organic matter transformation, bacterial carbon production, and diversity in a shallow oligotrophic Aquifer: Results From flow-through sediment microcosm experiments. Frontiers in Microbiology, 11. 10.3389/fmicb.2020.543567

Hose, G. C., Chariton, A., Daam, M. A., Di Lorenzo, T., Galassi, D. M. P., Halse, S. A., Reboleira, A. S. P. S., Robertson, A. L., Schmidt, S. I., & Korbel, K. L. (2022). Invertebrate traits, diversity and the vulnerability of groundwater ecosystems. Functional Ecology, 39(9), 2200–2214. 10.1111/1365-2435.14125

IPCC. (2022). Climate Change 2022: Impacts, Adaptation and Vulnerability. Contribution of Working Group II to the Sixth Assessment Report of the Intergovernmental Panel on Climate Change [H.-O. Pörtner, D.C. Roberts, M. Tignor, E.S. Poloczanska, K. Mintenbeck, A. Alegría, M. Craig, S. Langsdorf, S. Löschke, V. Möller, A. Okem, B. Rama (Eds.)]. Cambridge University Press. Cambridge University Press, Cambridge, UK and New York, NY, USA, 1–3056. 10.1017/9781009325844

Jones, A. A., & Bennett, P. C. (2014). Mineral Microniches Control the Diversity of Subsurface Microbial Populations. Geomicrobiology Journal, 31(3), 246–261. 10.1080/01490451.2013.809174

Karwautz, C., Zhou, Y., Kerros, M.-E., Weinbauer, M. G., & Griebler, C. (2022). Bottom-Up Control of the Groundwater Microbial Food-Web in an Alpine Aquifer. Frontiers in Ecology and Evolution, 10. https://www.frontiersin.org/article/10.3389/fevo.2022.854228

Kelland, M. E., Wade, P. W., Lewis, A. L., Taylor, L. L., Sarkar, B., Andrews, M. G., Lomas, M. R., Cotton, T. E. A., Kemp, S. J., James, R. H., Pearce, C. R., Hartley, S. E., Hodson, M. E., Leake, J. R., Banwart, S. A., & Beerling, D. J. (2020). Increased yield and CO2 sequestration potential with the C4 cereal Sorghum bicolor cultivated in basaltic rock dust-amended agricultural soil. Global Change Biology, 26(6), 3658–3676. 10.1111/gcb.15089

Kellermann, C., Selesi, D., Lee, N., Hügler, M., Esperschütz, J., Hartmann, A., & Griebler, C. (2012). Microbial CO2 fixation potential in a tar-oil-contaminated porous aquifer. FEMS Microbiology Ecology, 81(1), 172–187. 10.1111/j.1574-6941.2012.01359.x

Kump, L. R., Brantley, S. L., & Arthur, M. A. (2000). Chemical Weathering, Atmospheric CO2, and Climate. Annual Review of Earth and Planetary Sciences, 28(1), 611–667. 10.1146/annurev.earth.28.1.611

Lampert, W., & Sommer, U. (2007). Limnoecology.

Langer, J. A. F., Sharma, R., Schmidt, S. I., Bahrdt, S., Nam, B., Achterberg, E. P., Riebesell, U., Horn, H. G., Boersma, M., Thines, M., & Schwenk, K. (2017). Community barcoding reveals little effect of ocean acidification on the composition of coastal plankton communities: Evidence from a long-term mesocosm study in the Gullmar Fjord, Skagerrak. PLoS ONE, 12(4), 1–20. 10.1594/PANGAEA.864598

Li, G., Hartmann, J., Derry, L. A., West, A. J., You, C.-F., Long, X., Zhan, T., Li, L., Li, G., Qiu, W., Li, T., Liu, L., Chen, Y., Ji, J., Zhao, L., & Chen, J. (2016). Temperature dependence of basalt weathering. Earth and Planetary Science Letters, 443, 59–69. 10.1016/j.epsl.2016.03.015

Liu, H., Chen, M., Zhu, F., & Harrison, P. J. (2016). Effect of Diatom Silica Content on Copepod Grazing, Growth and Reproduction. Frontiers in Marine Science, 3. https://www.frontiersin.org/articles/10.3389/fmars.2016.00089

Macpherson, G. L., Roberts, J. A., Blair, J. M., Townsend, M. A., Fowle, D. A., & Beisner, K. R. (2008).Increasing shallow groundwater CO2 and limestone weathering, Konza Prairie, USA. Geochimica et Cosmochimica Acta, 72(23), 5581–5599. 10.1016/j.gca.2008.09.004

Macpherson, G. L., & Sullivan, P. L. (2019). Watershed-scale chemical weathering in a merokarst terrain, northeastern Kansas, USA. Chemical Geology, 527, 118988. 10.1016/j.chemgeo.2018.12.001

Maher, K. (2010). The dependence of chemical weathering rates on fluid residence time. Earth and Planetary Science Letters, 294(1–2), 101–110. 10.1016/j.epsl.2010.03.010

Malard, F., & Hervant, F. (1999). Oxygen supply and the adaptations of animals in groundwater. Freshwater Biology, 41(1), 1–30. 10.1046/j.1365-2427.1999.00379.x

Mammola, S., Cardoso, P., Culver, D. C., Deharveng, L., Ferreira, R. L., Fišer, C., Galassi, D. M. P., Griebler, C., Halse, S., Humphreys, W. F., Isaia, M., Malard, F., Martinez, A., Moldovan, O. T., Niemiller, M. L., Pavlek, M., Reboleira, A. S. P. S., Souza-Silva, M., Teeling, E. C., … Zagmajster, M. (2019).Scientists’ Warning on the Conservation of Subterranean Ecosystems. BioScience, 69(8), 641–650. 10.1093/biosci/biz064

Marçais, J., Gauvain, A., Labasque, T., Abbott, B. W., Pinay, G., Aquilina, L., Chabaux, F., Viville, D., & de Dreuzy, J.-R. (2018). Dating groundwater with dissolved silica and CFC concentrations in crystalline aquifers. Science of The Total Environment, 636, 260–272. 10.1016/j.scitotenv.2018.04.196

Marxsen, J., Rütz, N. K., & Schmidt, S. I. (2021). Organic carbon and nutrients drive prokaryote and metazoan communities in a floodplain aquifer. Basic and Applied Ecology, 51,43–58. 10.1016/j.baae.2020.12.006

Matzke, D., & Hahn, H. J. (2002). Vergleich der Grundwasserfauna in Lockergesteins- und in Kluftgrundwasserleitern unter vergleichender Anwendung unterschiedlicher Sammeltechniken. Final Report Project Az HA 3214/1-1.

Meckenstock, R. U., Elsner, M., Griebler, C., Lueders, T., Stumpp, C., Aamand, J., Agathos, S. N., Albrechtsen, H.-J., Bastiaens, L., Bjerg, P. L., Boon, N., Dejonghe, W., Huang, W. E., Schmidt, S. I., Smolders, E., Sørensen, S. R., Springael, D., & van Breukelen, B. M. (2015). Biodegradation: Updating the concepts of control for microbial cleanup in contaminated aquifers. Environmental Science & Technology, 49(12), 7073–7081. 10.1021/acs.est.5b00715

Meybeck, M. (1993). Riverine transport of atmospheric carbon: Sources, global typology and budget.Water, Air, and Soil Pollution, 70, 443–463.

Moore, C. M., Mills, M. M., Arrigo, K. R., Berman-Frank, I., Bopp, L., Boyd, P. W., Galbraith, E. D., Geider, R. J., Guieu, C., Jaccard, S. L., Jickells, T. D., La Roche, J., Lenton, T. M., Mahowald, N. M., Marañón, E., Marinov, I., Moore, J. K., Nakatsuka, T., Oschlies, A., … Ulloa, O. (2013). Processes and patterns of oceanic nutrient limitation. Nature Geoscience, 6(9), Article 9. 10.1038/ngeo1765

Négrel, P., Allègre, C. J., Dupré, B., & Lewin, E. (1993). Erosion sources determined by inversion of major and trace element ratios and strontium isotopic ratios in river water: The Congo Basin case. Earth and Planetary Science Letters, 120(1), 59–76. 10.1016/0012-821X(93)90023-3

Nydahl, A. C., Wallin, M. B., Laudon, H., & Weyhenmeyer, G. A. (2020). Groundwater carbon within a boreal catchment: Spatiotemporal variability of a hidden aquatic carbon pool. Journal of Geophysical Research: Biogeosciences, 125(1), e2019JG005244. 10.1029/2019JG005244

Oksanen, J., Simpson, G., Blanchet, F., Kindt, R., Legendre, P., Minchin, P., O’Hara, R., Solymos, P., Stevens, M., Szoecs, E., Wagner, H., Barbour, M., Bedward, M., Bolker, B., Borcard, D., Carvalho, G., Chirico, M., De Caceres, M., Durand, S., … Weedon, J. (2022). Vegan: Community Ecology Package. R package version 2.6-2 [Computer software]. https://CRAN.R-project.org/package=vegan

Overholt, W. A., Trumbore, S., Xu, X., Bornemann, T. L. V., Probst, A. J., Krüger, M., Herrmann, M., Thamdrup, B., Bristow, L. A., Taubert, M., Schwab, V. F., Hölzer, M., Marz, M., & Küsel, K. (2022). Carbon fixation rates in groundwater similar to those in oligotrophic marine systems. Nature Geoscience, 1–7. 10.1038/s41561-022-00968-5

Penman, D. E., Caves Rugenstein, J. K., Ibarra, D. E., & Winnick, M. J. (2020). Silicate weathering as a feedback and forcing in Earth’s climate and carbon cycle. Earth-Science Reviews, 209, 103298. 10.1016/j.earscirev.2020.103298

R Core Team. (2023). R: A Language and Environment for Statistical Computing [Computer software]. The R Foundation for Statistical Computing. https://www.R-project.org/

Ramaekers, L., Pinceel, T., Brendonck, L., & Vanschoenwinkel, B. (2022). Direct effects of elevated dissolved CO2 can alter the life history of freshwater zooplankton. Scientific Reports, 12(1), Article 1.10.1038/s41598-022-10094-2

Rau, G. C., Cuthbert, M. O., Andersen, M. S., Baker, A., Rutlidge, H., Markowska, M., Roshan, H., Marjo, C. E., Graham, P. W., & Acworth, R. I. (2015). Controls on cave drip water temperature and implications for speleothem-based paleoclimate reconstructions. Quaternary Science Reviews, 1–18. 10.1016/j.quascirev.2015.03.026

Raymond, P. A., & Cole, J. J. (2003). Increase in the Export of Alkalinity from North America’s Largest River. Science, 301(5629), 88–91. 10.1126/science.1083788

Rue, G. P., & McKnight, D. M. (2021). Enhanced rare earth element mobilization in a mountain watershed of the Colorado mineral belt with concomitant detection in aquatic biota: Increasing climate change-driven degradation to water quality. Environmental Science & Technology, 55(21), 14378–14388. 10.1021/acs.est.1c02958

Sabine, C. L., Feely, R. A., Gruber, N., Key, R. M., Lee, K., Bullister, J. L., Wanninkhof, R., Wong, C. S., Wallace, D. W. R., Tilbrook, B., Millero, F. J., Peng, T.-H., Kozyr, A., Ono, T., & Rios, A. F. (2004). The oceanic sink for anthropogenic CO2. Science, 305(5682), 367–371. 10.1126/science.1097403

Schmidt, S. I., Cuthbert, M. O., & Schwientek, M. (2017). Towards an integrated understanding of how micro scale processes shape groundwater ecosystem functions. Science of the Total Environment, 592, 215–227. 10.1016/j.scitotenv.2017.03.047

Schmidt, S. I., & Hahn, H. J. (2012). What is groundwater and what does this mean to fauna? – An opinion. Limnologica, 42(1), 1–6. 10.1016/j.limno.2011.08.002

Schmidt, S. I., Hahn, H. J., Watson, G. D., Woodbury, R. J., & Hatton, T. J. (2004). Sampling fauna in stream sediments as well as groundwater using one net sampler. Acta Hydrochimica et Hydrobiologica, 32(2), 131–137. 10.1002/aheh.200300522

Schmidt, S. I., Kodeš, V., Svátková, M., & Blabolil, P. (2021). Ekologie podzemních vod, po stopách Františka Vejdovského (Czech with English abstract: Groundwater ecology, on Frantisek Vejdovsky’s tracks). Vodní Hospodářství, 71(2), 13–19.

Schwartzman, D. W., & Volk, T. (1989). Biotic enhancement of weathering and the habitability of Earth. Nature, 340(6233), Article 6233. 10.1038/340457a0

Spengler, C., & Hahn, J. (2018). Thermostress: Ökologisch begründete, thermische Schwellenwerte und Bewertungsansätze für das Grundwasser. KW Korrespondenz Wasserwirtschaft, 11(9), 521–525. 10.3243/kwe2018.09.001

UNESCO World Water Assessment Programme. (2022). The United Nations World Water Development Report 2022: Groundwater: Making the invisible visible [UNESCO Digital Library]. https://unesdoc.unesco.org/ark:/48223/pf0000380721

Vejvodová, J., Hušková, R., Sklenář, M., Nesměráková, E., Růžička, P., Schinkmanová, H., & Pospíšilová, J. (2006). B2.2 Drinking Water. In Yearbook Prague Environment 2006. https://envis.praha.eu/rocenky/Pr06_htm/aB2_02.htm#B2_021

Vicca, S., Goll, D. S., Hagens, M., Hartmann, J., Janssens, I. A., Neubeck, A., Peñuelas, J., Poblador, S., Rijnders, J., Sardans, J., Struyf, E., Swoboda, P., van Groenigen, J. W., Vienne, A., & Verbruggen, E. (2022). Is the climate change mitigation effect of enhanced silicate weathering governed by biological processes? Global Change Biology, 28(3), 711–726. 10.1111/gcb.15993

Walker, J. C. G., Hays, P. B., & Kasting, J. F. (1981). A negative feedback mechanism for the long-term stabilization of Earth’s surface temperature. Journal of Geophysical Research, 86(C10), 9776. 10.1029/JC086iC10p09776

Weiss, L. C., Pötter, L., Steiger, A., Kruppert, S., Frost, U., & Tollrian, R. (2018). Rising pCO2 in freshwater ecosystems has the potential to negatively affect predator-induced defenses in Daphnia. Current Biology, 28(2), 327–332.e3. 10.1016/j.cub.2017.12.022

Zahradníček, P., Brázdil, R., štěpánek, P., & Trnka, M. (2021). Reflections of global warming in trends of temperature characteristics in the Czech Republic, 1961–2019. International Journal of Climatology, 41(2), 1211–1229. 10.1002/joc.6791

Zhang, S., Bai, X., Zhao, C., Tan, Q., Luo, G., Wang, J., Li, Q., Wu, L., Chen, F., Li, C., Deng, Y., Yang, Y., & Xi, H. (2021). Global CO2 consumption by silicate rock chemical weathering: Its past and future. Earth’s Future, 9(5), e2020EF001938. 10.1029/2020EF001938

