## Supplementary Information for "Direct and indirect effects of the increase in atmospheric CO_2_ and temperature on groundwater organisms"

*Table S1: Sampling sites and information (separate file)*

*Table S2: Fauna occurrences in the wells (separate file)*

*Table S3: Parameter of linear models for trends in chemical and physical properties over time (separate file)*

### Supplementary Information (SI) 1 - Summary of $CO_2$-driven weathering

In general, chemical weathering of silicate minerals "consumes" (Hartmann et al., 2009; Walker et al., 1981) atmospheric $CO_2$ (e.g. eq. 1, Meybeck, 1993). Depending on the prevalent pH, the $CO_2$ is transformed into dissolved bicarbonate, the negative charge of which is balanced by cations released from chemical weathering of minerals (Moosdorf et al., 2011); e.g. eq. 1, from Meybeck (1993); for mineral-specific equations, refer to e.g. Marx et al. (2017). Dissolved silica ($SiO_2$) is formed, or $SiO_4^{4-}$, or $Si(OH)_4$, or orthosilicate, i.e. $SiO_3^{2-}$, depending e.g. on background pH, hydration, formation of secondary minerals, etc. (Bailey, 1991; Bakalowicz, 1994; Blume et al., 2010)).

Eq 1. Non-carbonate mineral + $CO_2$ + $H_20$ → $HCO_3^-$ + clay mineral + cation + $SiO_2$

The consumed $CO_2$ is transported by rivers (Meybeck, 1993) and groundwater (Calmels et al., 2011; Luijendijk et al., 2020) to the oceans in the form of $HCO_3^-$. There, the $HCO_3^-$ precipitates to form carbonates (calcite and dolomite (Zhang et al., 2021; Oliva et al., 2003; Shaughnessy et al., 2021). This sink is called the Silicate Carbon Sink (SCS), in contrast to the Carbonate rock weathering Carbon Sink (CCS; Zhang et al., 2021). The area that the SCS covers, is 37.2 million $km^2$ larger than that covered by the CCS (Xiong et al., 2022) and thus potentially more influential on global patterns. On geological time scales, when the climate cools, the silicate weathering rate, replenishing the silicate carbon sink, slows down. This causes $CO_2$ concentrations to increase. This in turns warms the surface (greenhouse effect; (Kasting, 2019). Such global warming promotes rock weathering, by which more $CO_2$ is absorbed (Penman et al., 2020; Zhang et al., 2021). The Earth's surface temperature is thus stabilized on a large scale by a negative feedback between atmospheric $CO_2$ and silicate weathering. This is why many researchers such as Brantley et al. (2023) and Li et al. (2016) call this feedback of silicate weathering on temperature the 'geological thermostat'. Without this thermostat and e.g. the ocean carbon sink (Lehmann et al., 2023), temperatures might already have risen more than they have, and might have turned the planet already uninhabitable (Brantley et al., 2023).

Independently of trends on geological time scales, the Earth's mean surface temperature has recently risen to an extent that raises concern. The main cause for the increase is seen in the greenhouse gases which result mainly from coal and oil burning (IPCC, 2022). With silicate weathering rates being strongly temperature-dependent (Dessert et al., 2003; Walker et al., 1981; Webb and Waling, 1992, White et al., 1999), the recently increased temperatures enhance silicate weathering, and thus as a net result, the consumption of atmospheric $CO_2$ (Moosdorf et al., 2011; Oliva et al., 2003; Vinnarasi et al., 2021; Walker et al., 1981). This may influence alkalinity through the increase in $HCO_3^-$. Since alkalinity is a major characteristic of the carbon cycle, (Lehmann et al., 2023) assume shorter term influences of the silicate weathering on the carbon cycle. The increasing atmospheric $CO_2$ concentrations that are measured are the net effect of an increase from anthropogenic sources and the counteracting silicate thermostat which stabilizes temperature on earth. Some researchers argue that the impact of climate on silicate weathering and vice versa is only relevant on geological timescales (Kasting, 2019; Walker et al., 1981). However, others show that silicate weathering responded recently rapidly to climate change (< 100 years; Beaulieu et al., 2012; Gislason et al., 2009). Accordingly, the silicate rock weathering carbon sink flux (SCSF) has risen recently (Zhang et al., 2021), and so have the stocks (Xiong et al., 2022).

In most of the research on silicate weathering and $CO_2$ consumption, only surficial weathering and surface runoff were considered. However, river flow may originate to a large fraction from groundwater which is embedded in weatherable material. In the rivers' catchments, precipitation infiltrates into the subsurface and follows gravity, mostly towards rivers, lakes, and oceans (Fitts, 2012). The flow paths through pores and fractures are often hydrogeologically complex (Calmels et al., 2011; Schmidt et al., 2017). During this passage, the water takes on a chemical signature characteristic of the parent material, as a net effect of the biogeochemical reactions having taken place (Li et al., 2021). This includes uptake of soil $CO_2$ (Bakalowicz, 1994). An increase of soil $CO_2$ was recently observed and attributed to increased plant productivity and increased soil microbiological respiration in response to increased atmospheric $CO_2$ (Andrews and Schlesinger, 2001). Thus, groundwater may increase in $CO_2$, although most groundwater aquifers are already super saturated with $CO_2$ (Macpherson, 2009). E.g., 20% $CO_2$ increase was observed in a limestone aquifer (Macpherson et al., 2008). In contrast to the surface lithologies which are prone to increased weathering directly from increased atmospheric $CO_2$ concentrations, the $CO_2$-triggered increase in groundwater silicate weathering is thus indirect. Depending on groundwater discharge rates, groundwater was estimated to contribute between 32% and 351% that from surficial silicate weathering (Zhang and Planavsky, 2019) and these authors estimate that groundwaters harbour 20% to 250% of chemical silicate weathering globally (Zhang & Planavsky, 2019a).

In most groundwater, increased temperatures might be another main driver  and in this case a direct driver , for weathering (Calmels et al., 2011). Because of the sheer surface area of lithologies exposed to subsurface weathering, groundwater can be expected to be a major contributor to the geological thermostat. Indeed, Zhang and Planavsky (2019) estimated that subsurface to surface silicate weathering might take up a proportion between 32% and 351% (Zhang and Planavsky, 2019). Processes in the subsurface, and in groundwater in particular, can thus be assumed to be at places more important for understanding global patterns of silicate weathering and its (indirect) effect on atmospheric $CO_2$ concentrations, than surface runoff-driven processes.

### Supplementary Information (SI) 2 – Methods - Map of the investigated area

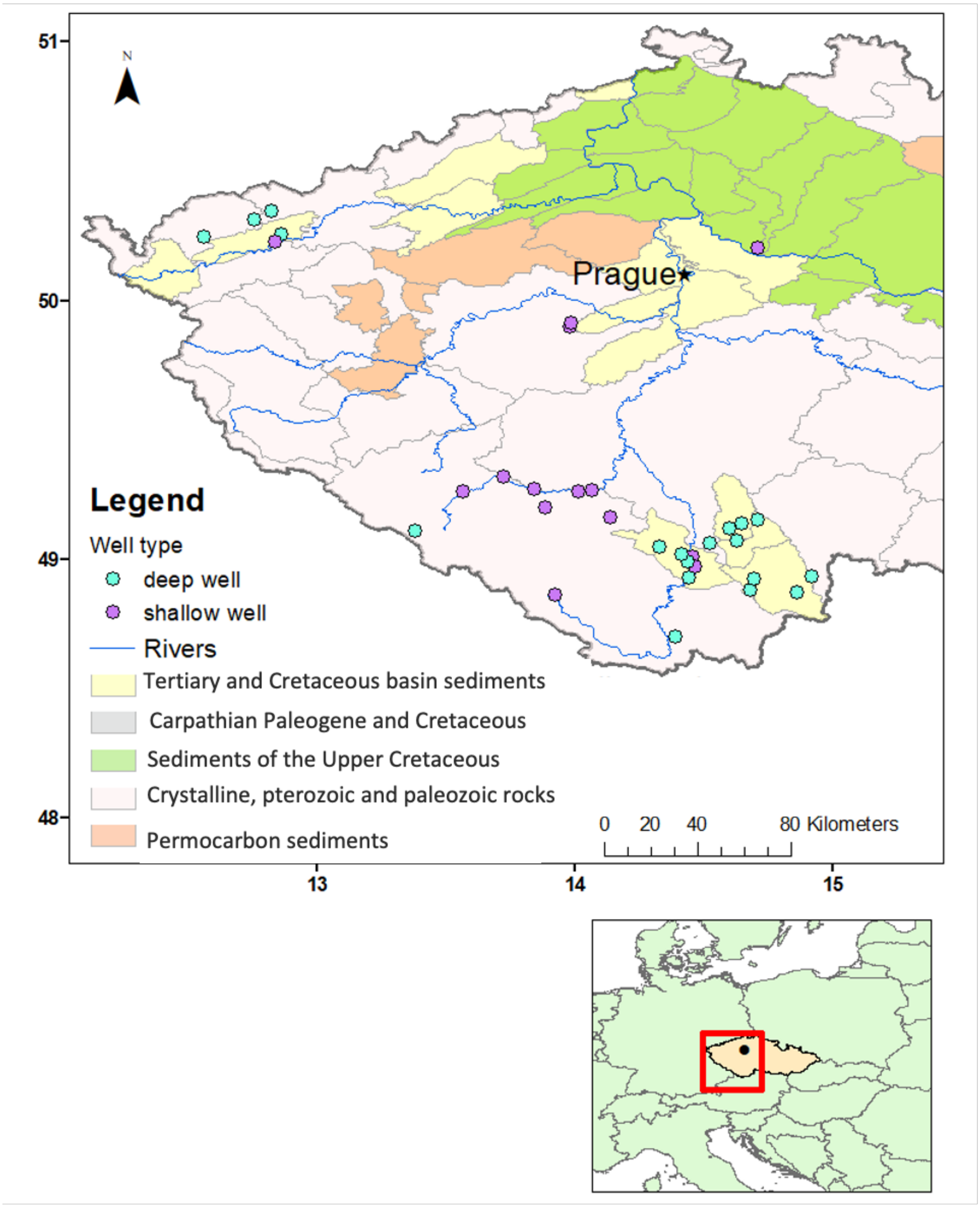

*Figure S1: Map of the sampling sites (top) and the situation of the sampled area within Europe (bottom). Coordinates shown in decimal degrees, WGS84.*

### Supplementary Information (SI) 3 – Methods - Further information on sampling, deriving, and evaluating $CO_2$

$CO_2$ in water, i.e. $CO_2$ (aq), hydrates and transforms to $H_2CO_3$. $H_2CO_3$ usually only makes up 1% of that of dissolved $CO_2$. The species cannot be distinguished, and their sum is often written as $H_2CO_3$* (Bailey, 1991). $H_2CO_3$* is often called free dioxide (Kopáček and Hezlar, 2021; Legler et al., 1986). Values for free dioxide were given in the database by CHMI only until 2011 included. Laboratory $CO_2$ measurements are biased by processes during sampling, storage and analysis, because $CO_2$ in water is transient (Bakalowicz, 2005). Usually, $CO_2$ (aq) and $pCO_2$ are thus derived via calculations (Macpherson et al., 2008). They depend on pH, gas pressure of other gases, density, species distribution, and total inorganic carbon. Not all of these influences were available in the database from CHMI. Using the R package "seacarb" v. 3.3.1 (Gattuso et al., 2022), pressure and atmospheric pressure were kept at the default values, since neither were available in the database. This is in both cases a source of error, since the pressure in groundwater is often above atmospheric pressure. However, "the small deviations from 1 atm total pressure which are encountered in many natural conditions may be neglected" (Weiss, 1974). Dissolved $CO_2$, however, depends (even more) strongly on the mixture of gases in solution and no other gas data were available except for $O_2$. The calculated dissolved $CO_2$ values have thus to be treated with care and we analysed in parallel $HCO_3^-$ with which $CO_2$ is in pH-dependent equilibrium and which was available for almost all samples.

### Supplementary Information (SI) 4 – Methods - Sampling of fauna

Since the debate on whether net samples or pumped samples are more representative of the fauna in the aquifers is on-going (Boulton et al., 2004; Hahn, 2003; Matzke and Hahn, 2002; Schmidt et al., 2004), it was checked whether total individual numbers or relative composition differed between samples from pumped water by adonis tests in the vegan R package (Oksanen et al., 2022). The volume for the net sampling was estimated based on the volume that the net might have passed through, with V = (radius of the net)$^2$ * $\pi$ * depth of the well, with the inner radius of the net being 0.025 m. Faunal individual numbers and biomass calculated from individual numbers using the dry masses in Marxsen et al. (2021) were not different in net-sampled versus in pumped samples (Figure S2 and Figure S3).

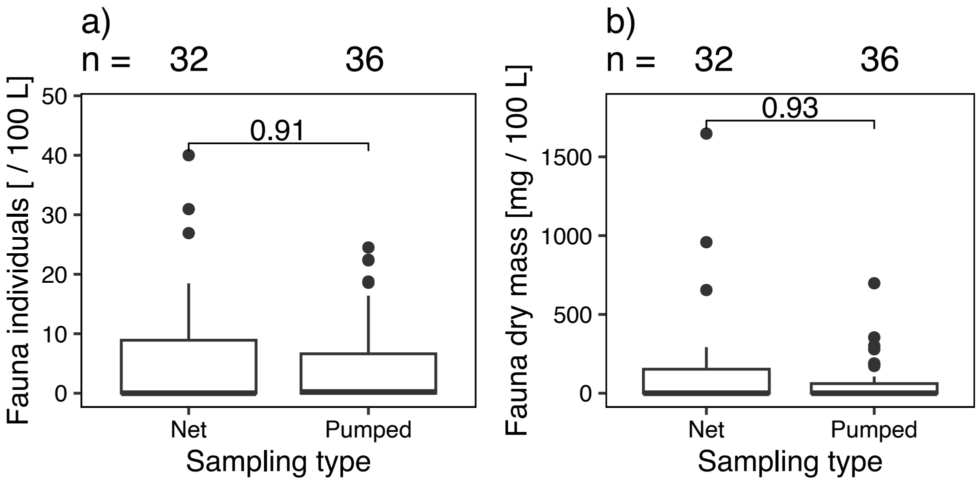

*Figure S2: Boxplots of the faunal individuals (a) and biomass (b) in net-sampled versus in pumped samples. The differences were not significant (Wilcoxon test), as shown by the included p levels.*

Faunal assemblages were not systematically different between net and pumped samples. This is visible in the adonis test in the vegan R package (Oksanen et al., 2022) which resulted in $R^2 = 0.16$, $p = 0.002$. Although the probability level was significant, the effect was much lower than 0.5, i.e. arbitrary. The MDS, where open circles signify net samples, and filled triangles, pumped samples (Figure S3), indicated a non-systematic difference between pumped and net samples.

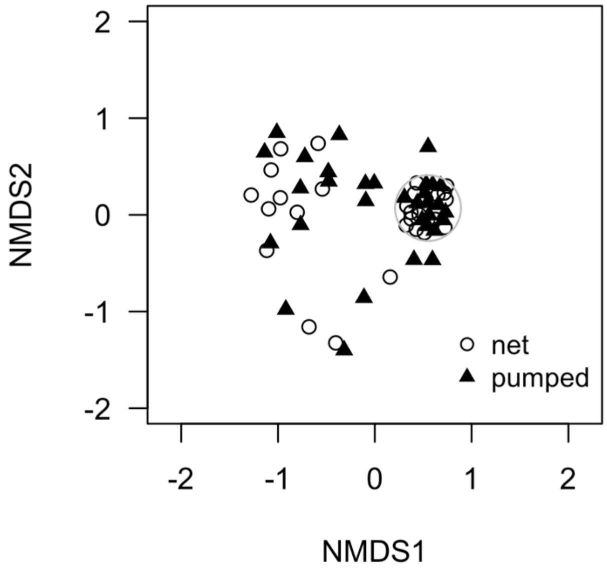

*Figure S3: Nonmetric MDS (NMDS) of the faunal assemblages in the wells. Symbols indicate the method. All samples surrounded by the grey circles were empty, thus fell into one point, and were drawn apart for clarification.*

### Supplementary Information (SI) 5 – Results - Chemical and physical properties over time in the investigated wells

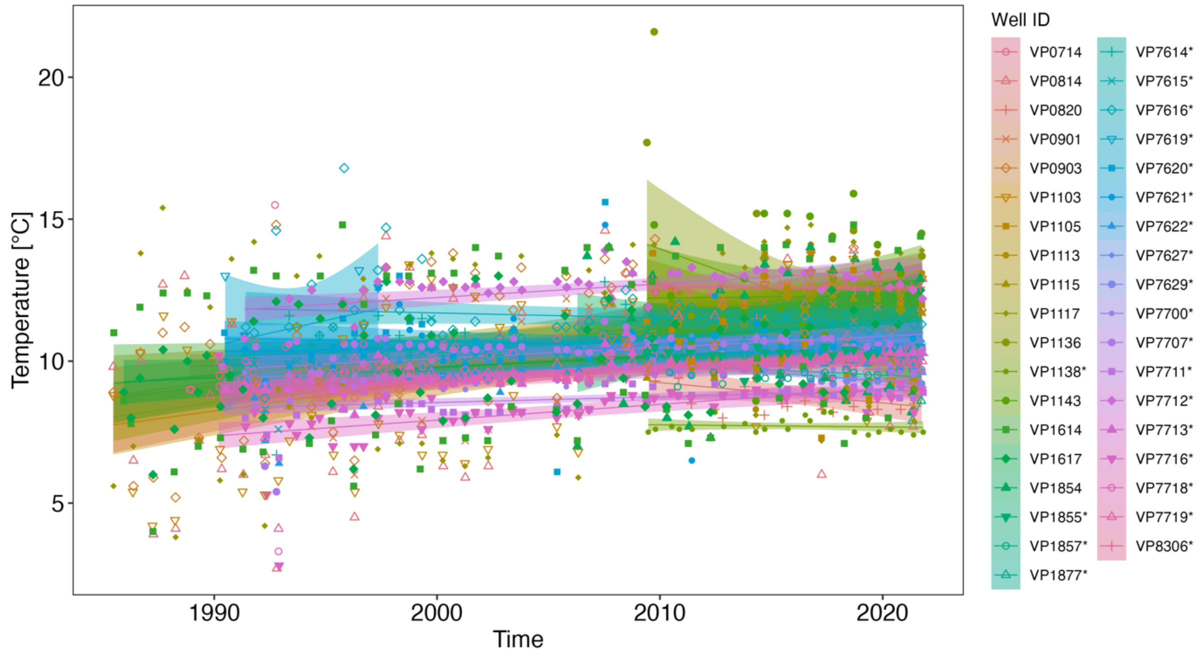

*Figure S4: Temperature over time in the wells that were sampled for fauna. Measurements, linear trend, and confidence margin are coloured according to the well. Well IDs of deep wells are marked with a star.*

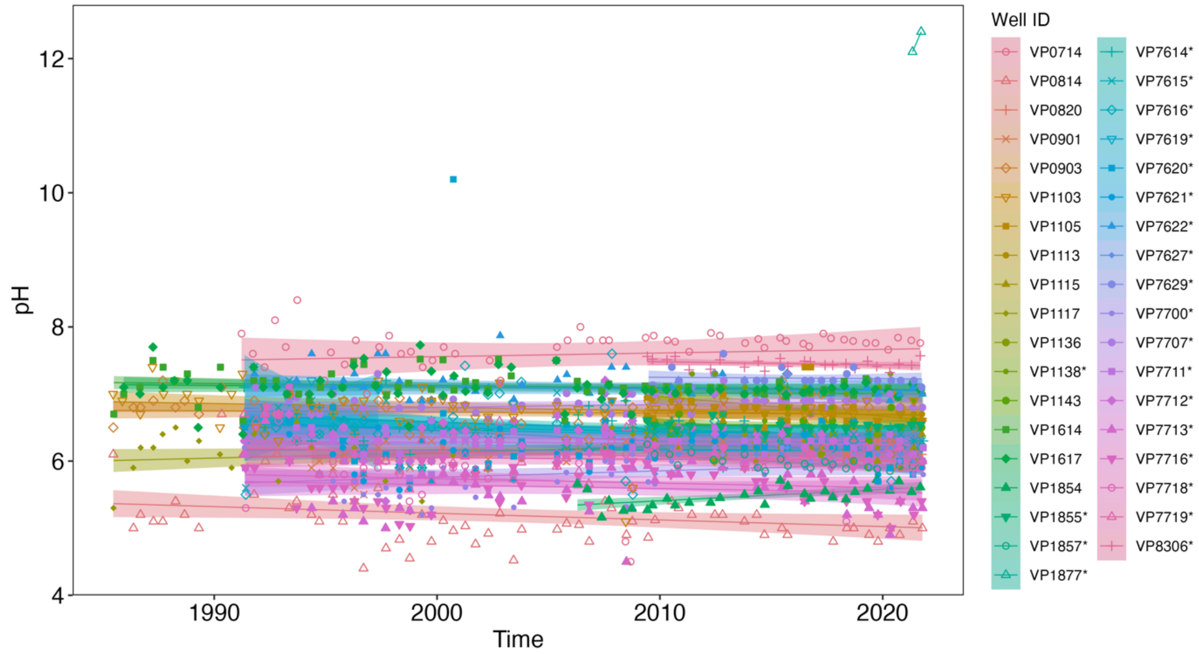

*Figure S5: pH over time in the wells that were sampled for fauna. Measurements, linear trend, and confidence margin are coloured according to the well. Well IDs of deep wells are marked with a star.*

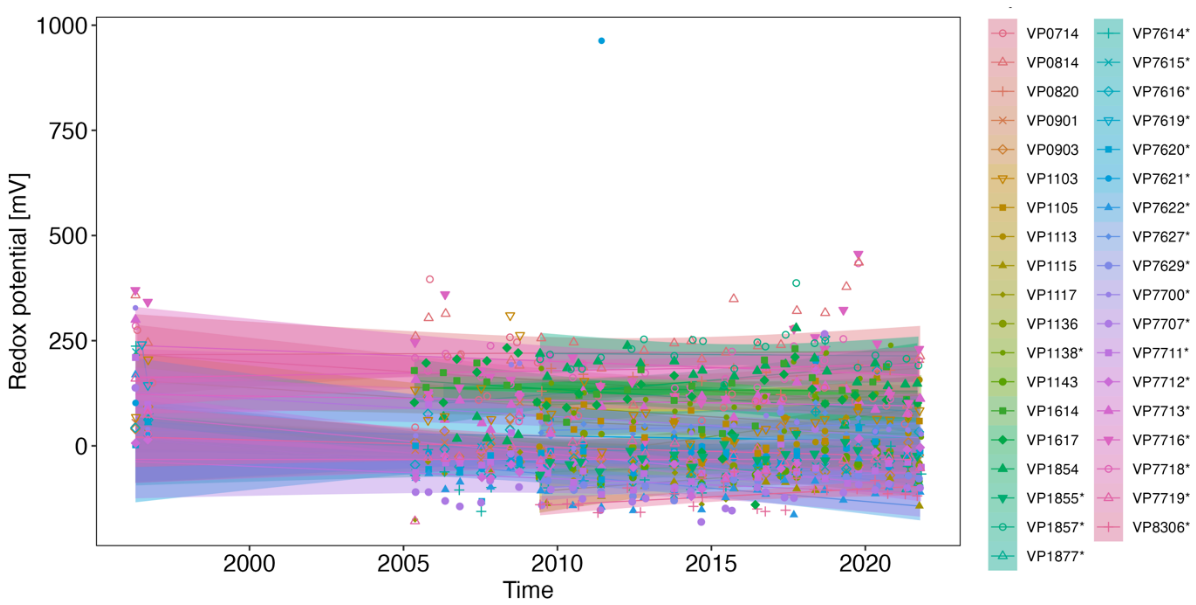

*Figure S6: Redox potential over time in the wells that were sampled for fauna. Measurements, linear trend, and confidence margin are coloured according to the well. Well IDs of deep wells are marked with a star.*

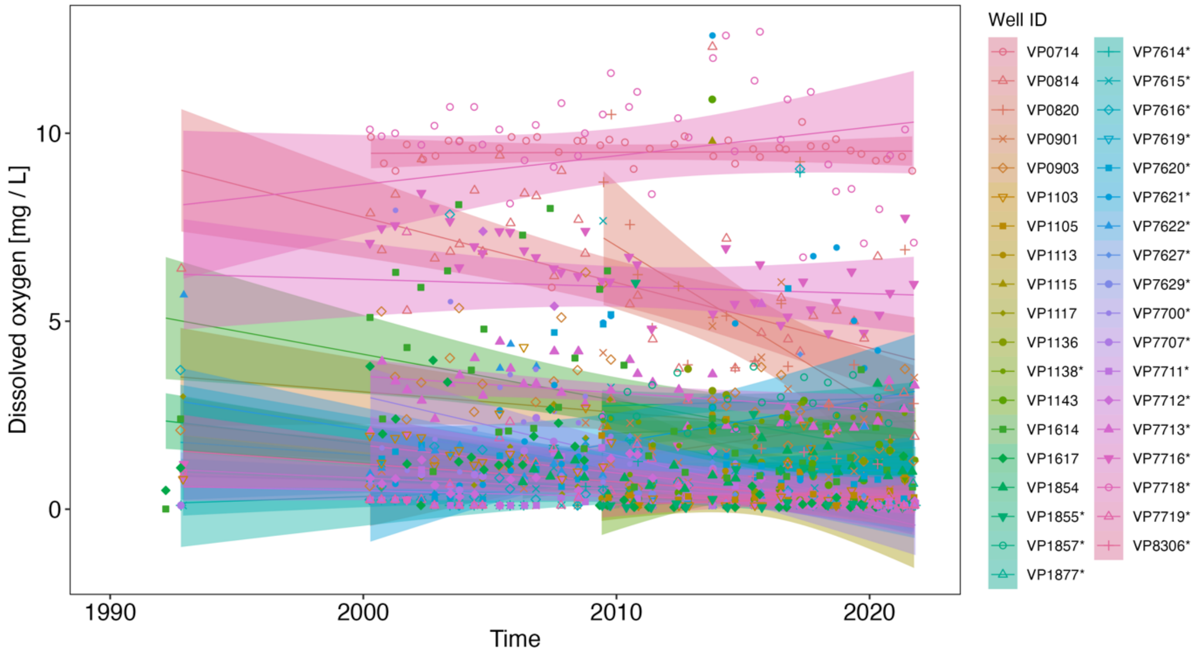

*Figure S7: Dissolved oxygen over time in the wells that were sampled for fauna. Measurements, linear trend, and confidence margin are coloured according to the well. Well IDs of deep wells are marked with a star.*

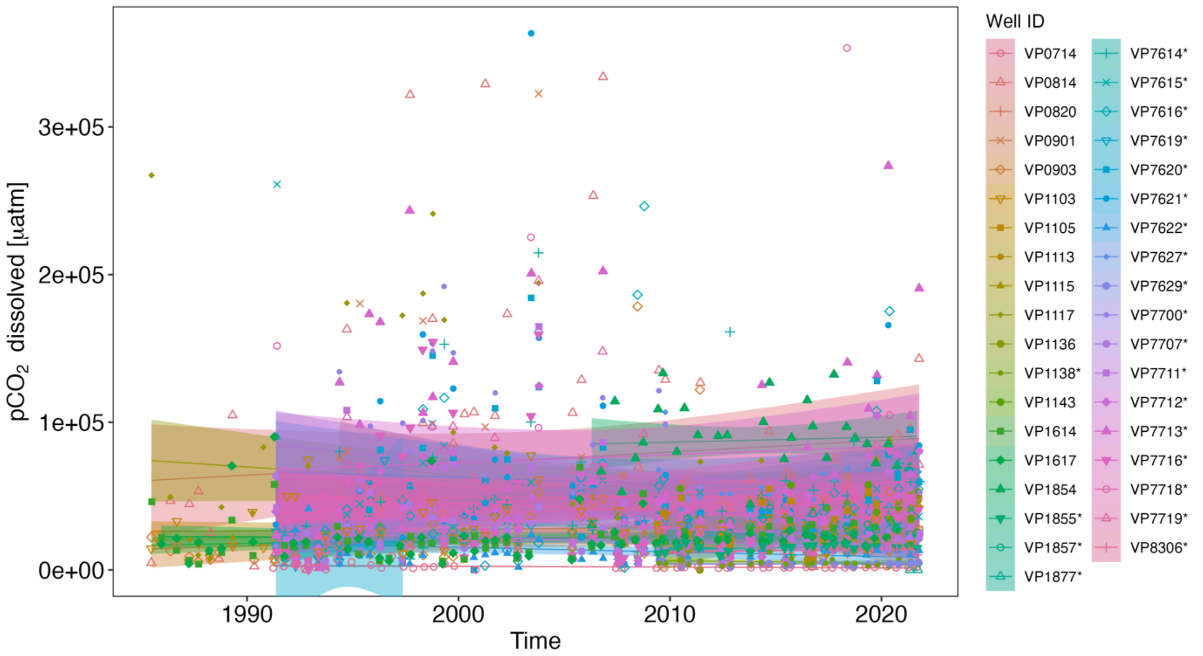

*Figure S8: Dissolved $pCO_2$ over time in the wells that were sampled for fauna. Measurements, linear trend, and confidence margin are coloured according to the well. Outliers above 400000 µatm were excluded for visualization purposes. Well IDs of deep wells are marked with a star.*

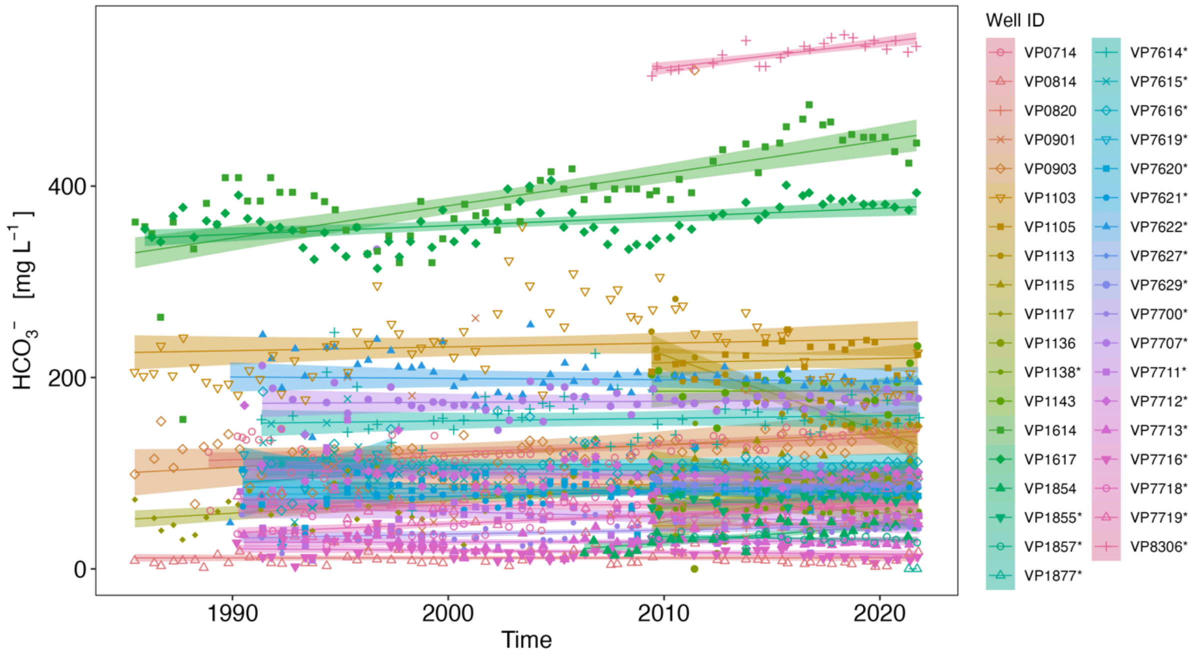

*Figure S9: $HCO_3^-$ concentrations over time in the wells that were sampled for fauna. Measurements, linear trend, and confidence margin are coloured according to the well. Well IDs of deep wells are marked with a star.*

### Supplementary Information (SI) 6 – Results - Molar ratios in the samples for classification into silicate versus carbonate weathering

Molar Ca/Na ratios in dependence on Mg/Na ratios can be used to identify whether waters have rather a carbonate weathering or a silicate weathering signature (Karlović et al., 2021). Except for a two samples, all values were below 1 for Ca/Na and Mg/Na, and thus reflected silicate.

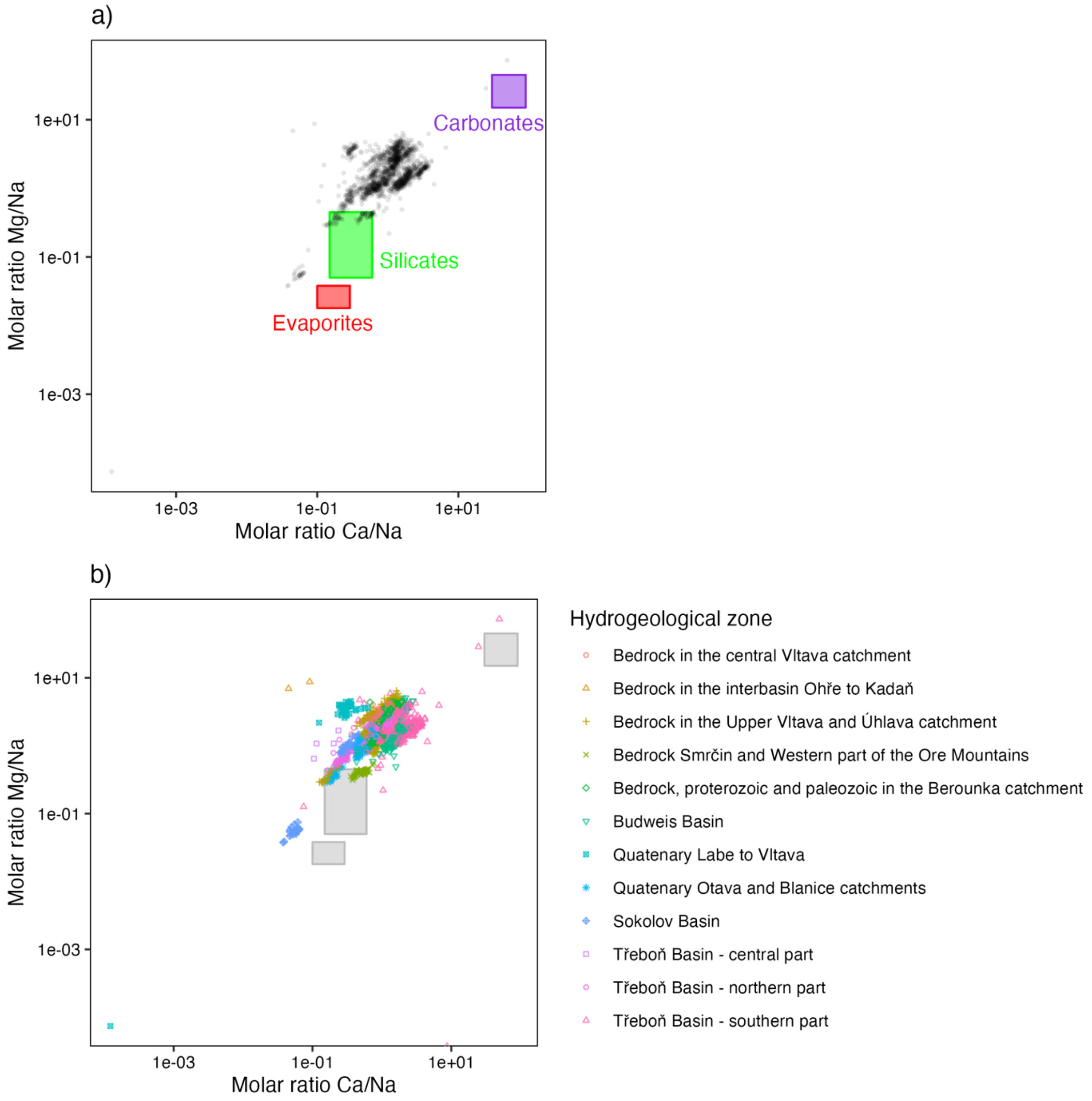

*Figure S10: Relationship between molar ratios of Ca and Na (x axis) and Mg and Na (y axis) in the well samples. The boxes indicate typical ranges of molar ratios according to Négrel et al. (1993) and Gaillardet et al. (1999). a) Points are transparent to indicate point densities, where there are overlays. b) Points are coloured according to the CHMI hydrogeological zone. Data from 1984 to 2021.*

### Supplementary Information (SI) 7 – Results - Fauna occurrence - map

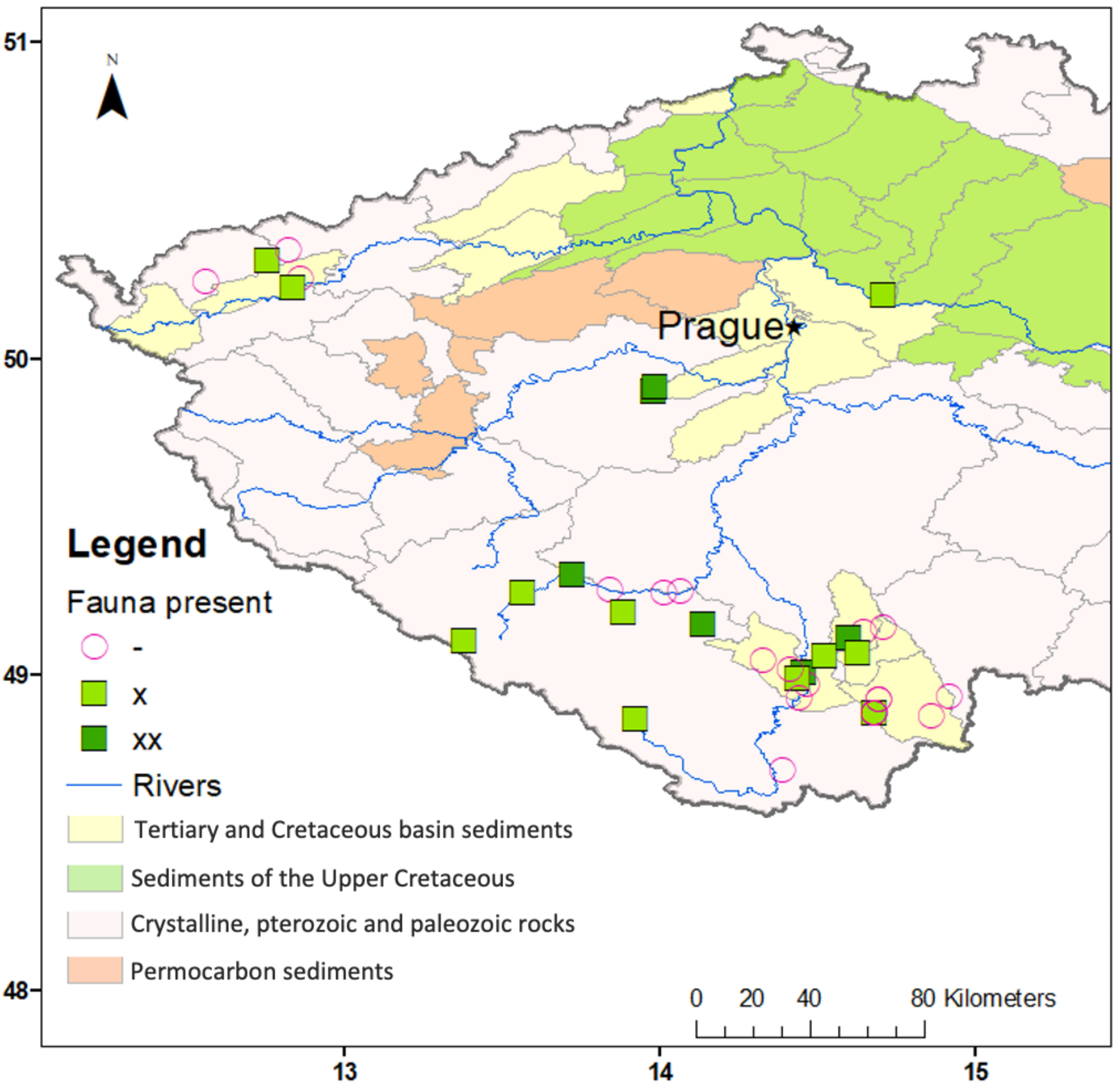

*Figure S11: Map of the spatial distribution of wells where high (xx), low (x) or no abundances (-) were found. The spatial distribution was not significant according to the Moran's I (see main text). The background of the hydrogeological zones is given in order to show that there also was no relationship with hydrogeological province. This was also tested statistically (see main text). Coordinates shown in decimal degrees, WGS84.*

### Supplementary Information (SI) 8 – Results - Fauna individual numbers, fauna dry mass, and prokaryote numbers in dependence on aquifer type

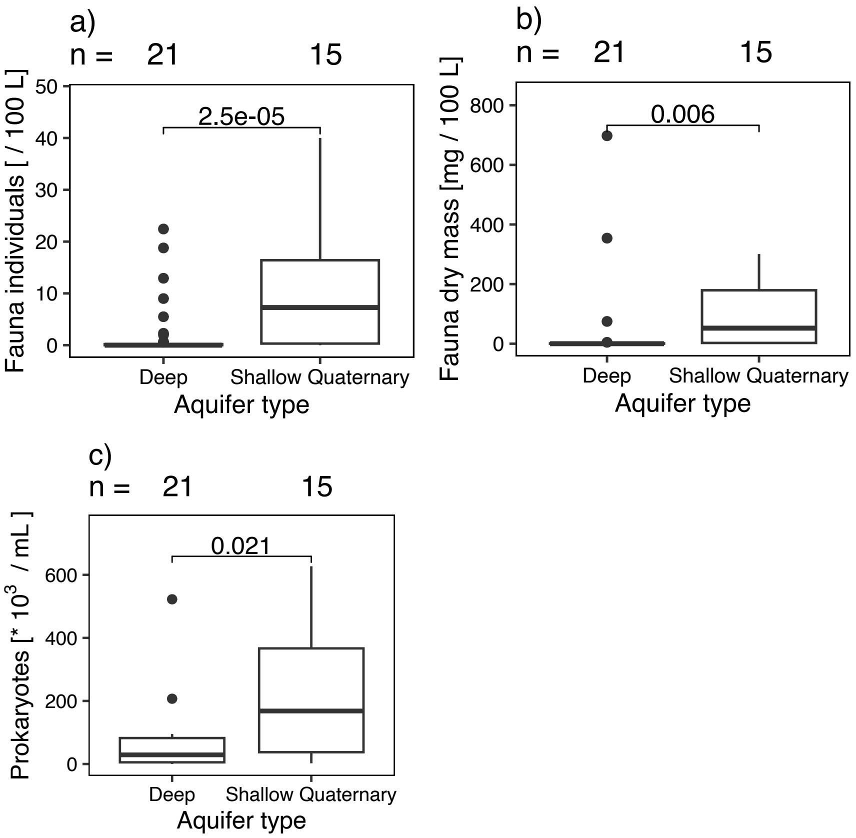

*Figure S12: Fauna individuals (a), biomass (b), and prokaryote cells (c) per aquifer type. The p-value of a Wilcoxon test is given in the respective figure.*

### Supplementary Information (SI) 9 – Results - Fauna individual numbers, fauna dry mass, and prokaryote numbers in dependence on each other and on chemical and physical properties

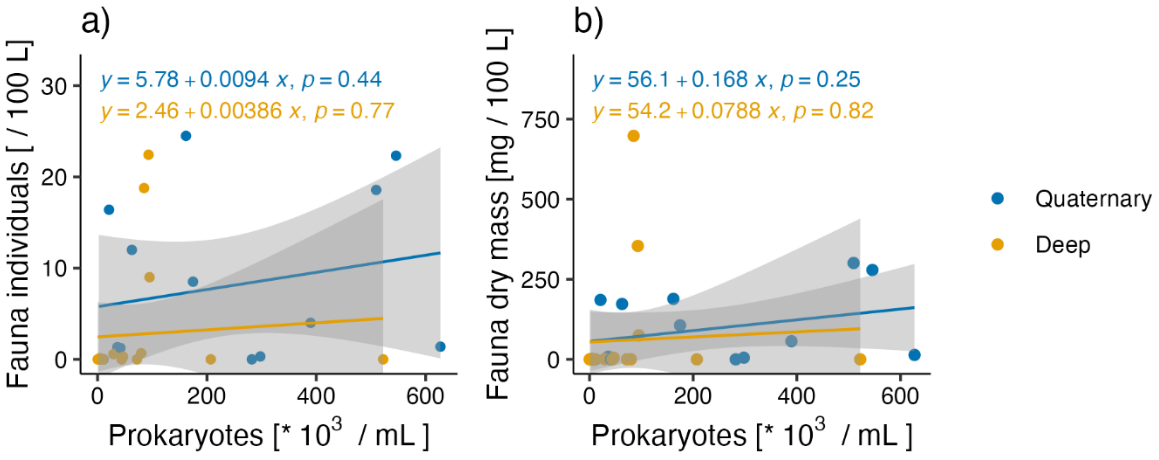

*Figure S13: Fauna abundances (a) and fauna biomass (b) in dependence on prokaryote numbers, separate fits per aquifer type. For the total relationships, without separation into aquifer type, and for the legend, see Figure 2 in the main paper.*

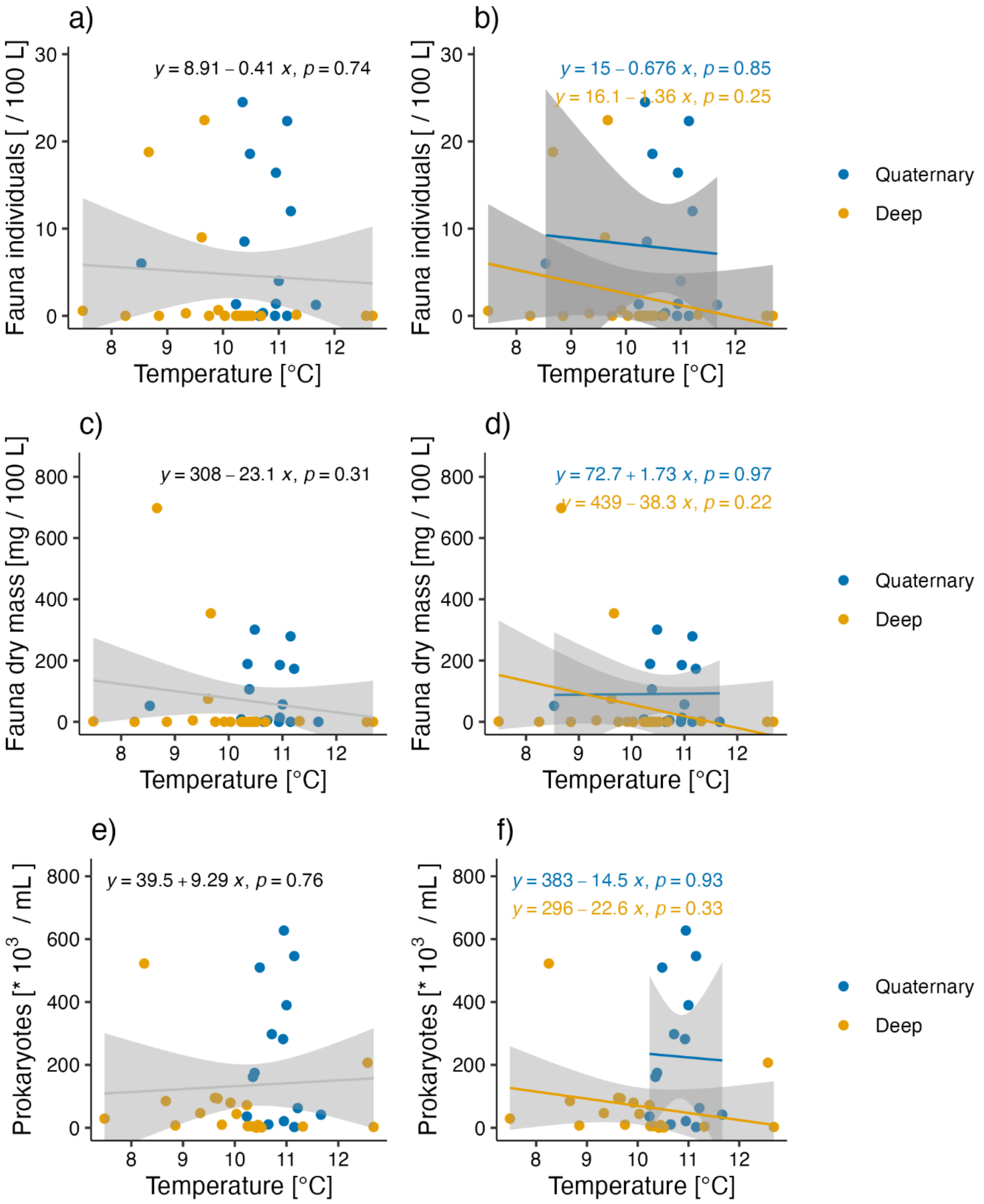

*Figure S14: Fauna individuals (a, b), dry mass (c, d), and prokaryote numbers (e, f) in dependence on temperature. In )b d), and f), linear models are shown separately for shallow (quaternary) and deep aquifers.*

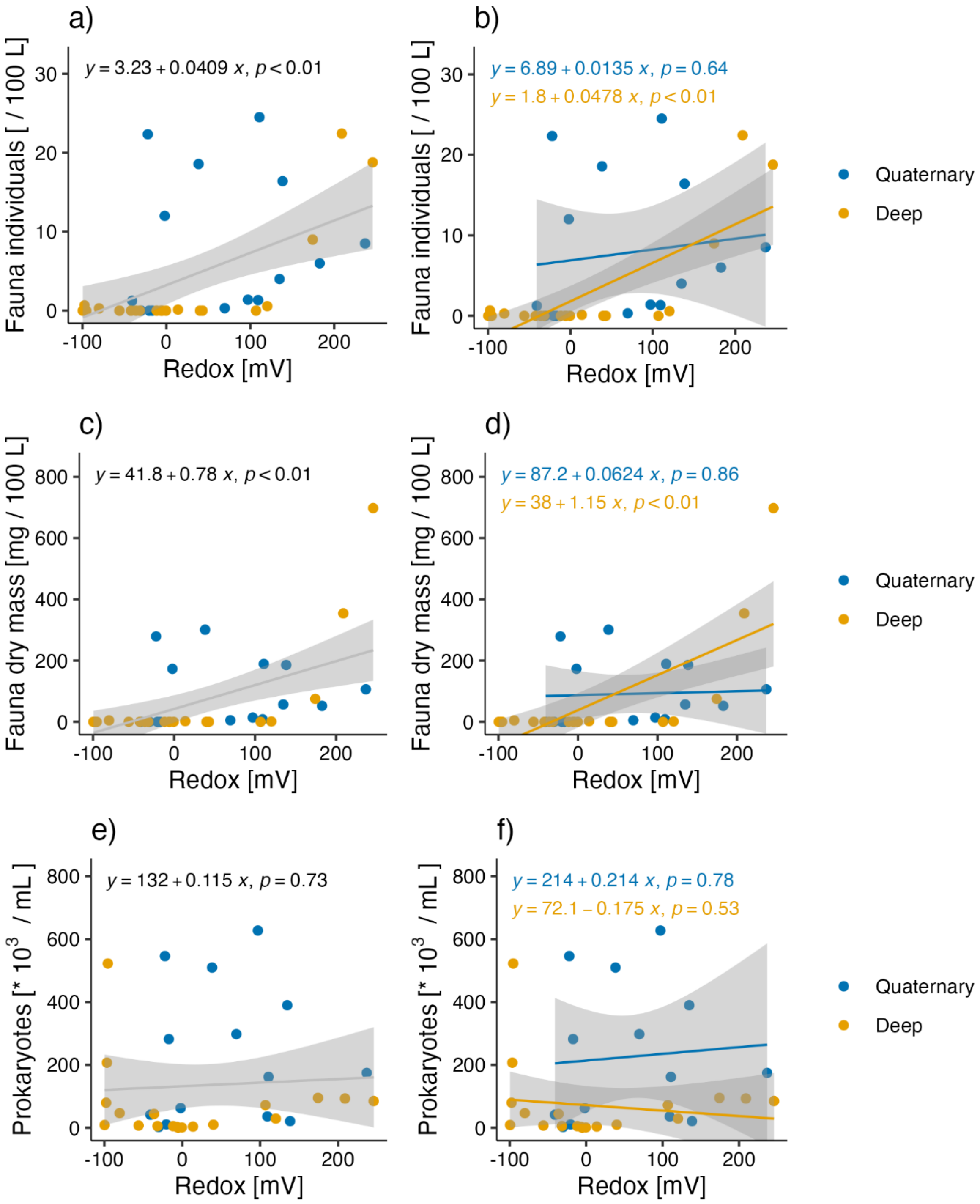

*Figure S15: Fauna individuals (a, b), fauna dry mass (c, d), and prokaryote numbers (e, f) in dependence on the redox potential. In b), d), and f), linear models are shown separately for shallow (quaternary) and deep aquifers.*

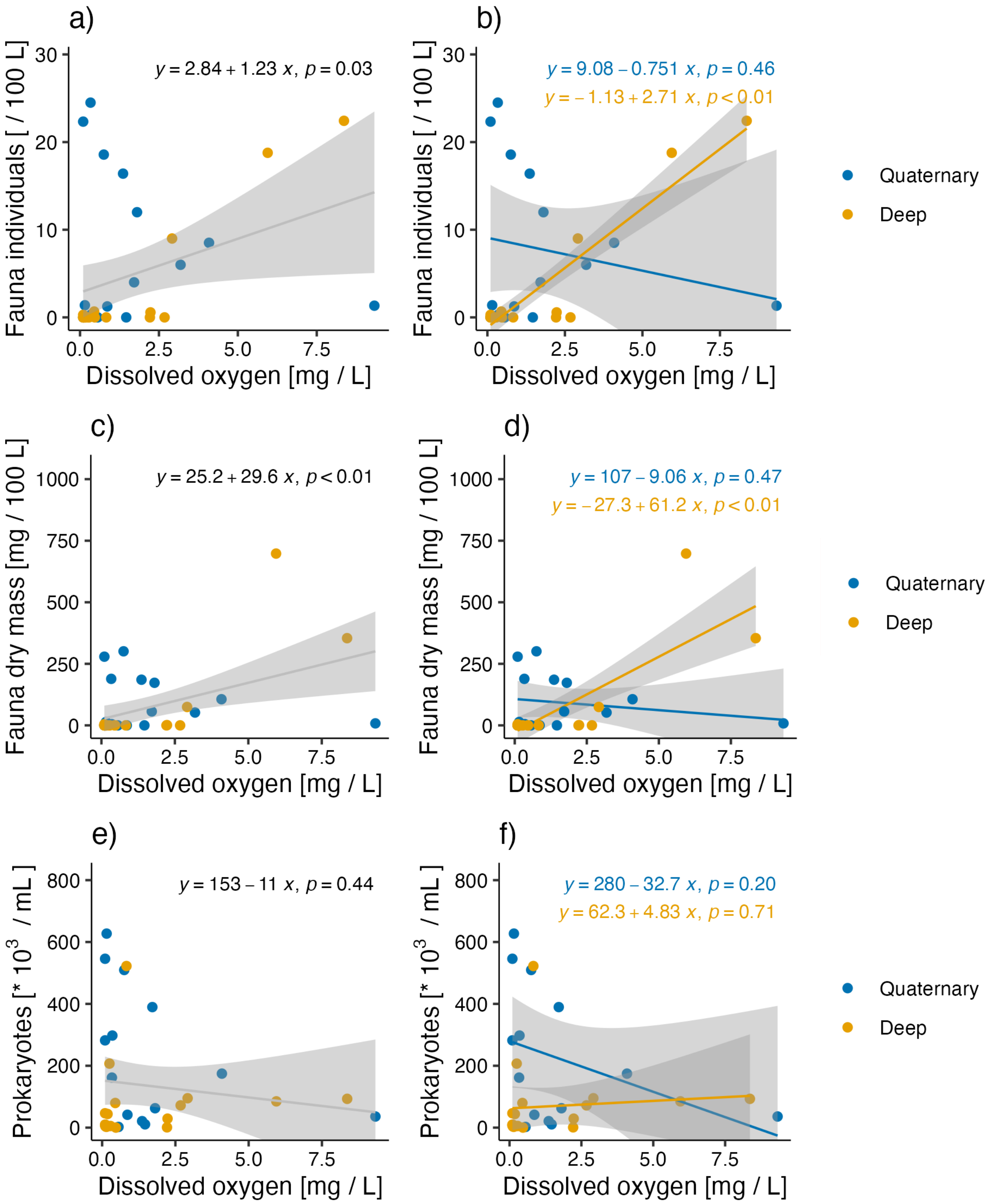

*Figure S16: Fauna individuals (a, b), fauna dry mass (c, d), and prokaryote numbers (e, f) in dependence on dissolved oxygen. In b), d), and f), linear models are shown separately for shallow (quaternary) and deep aquifers.*

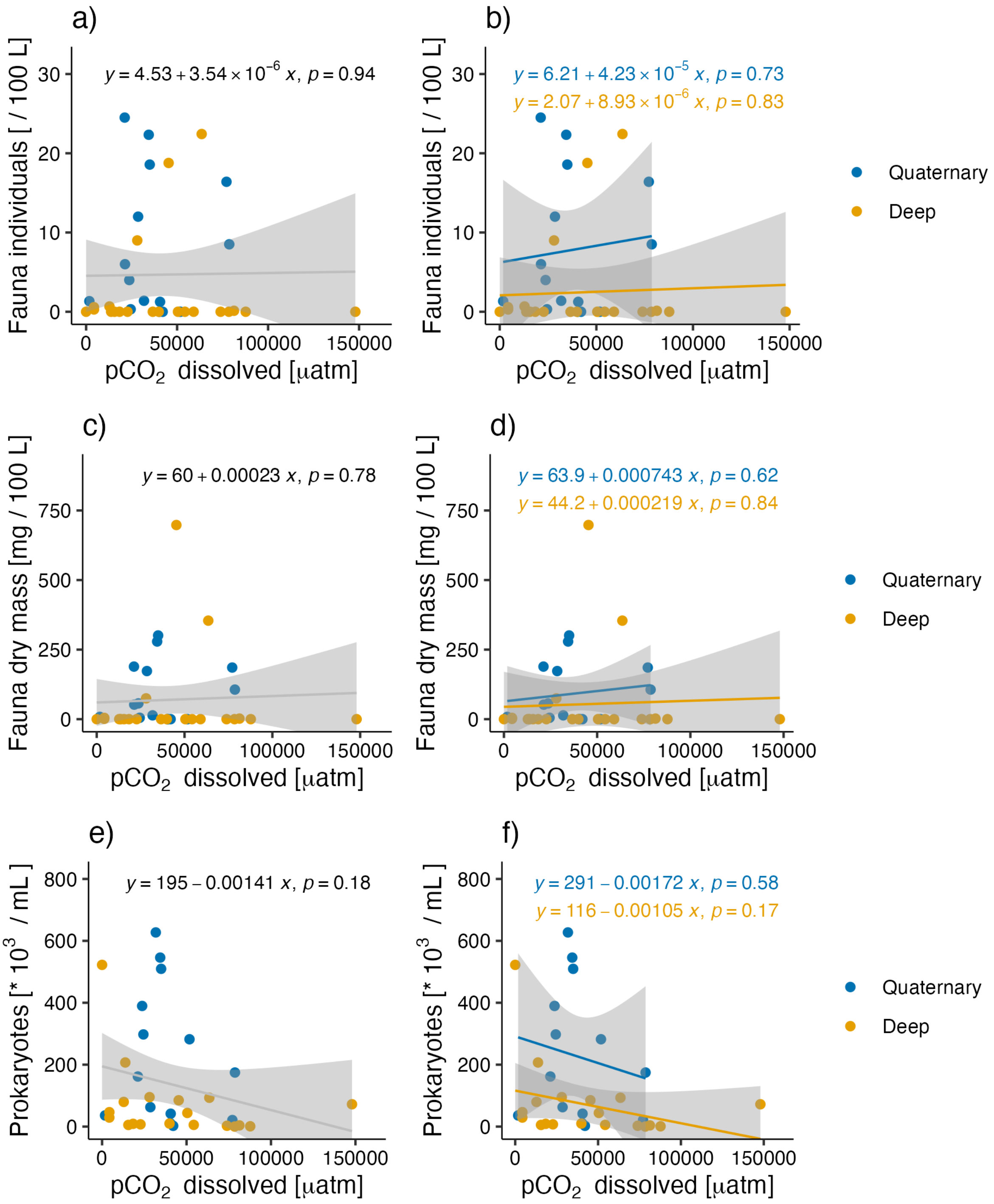

*Figure S17: Fauna individuals (a, b), fauna dry mass (c, d), and prokaryote numbers (e, f) in dependence on the concentration of $pCO_2$. In b), d), and f), linear models are shown separately for shallow (quaternary) and deep aquifers.*

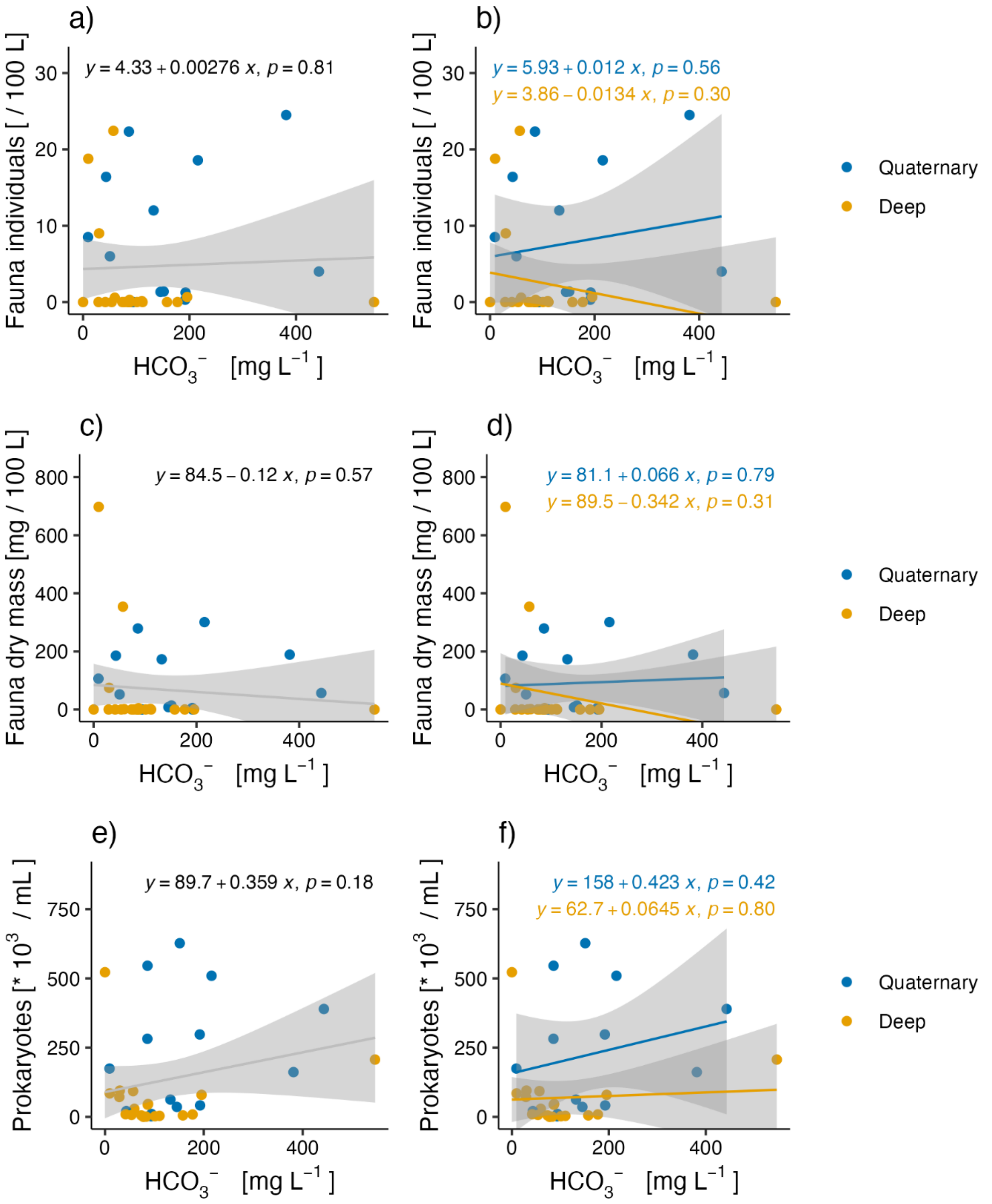

*Figure S18: Fauna individuals (a, b), dry mass (c, d), and prokaryote numbers (e, f) in HCO3. In b), d), and f), linear models are shown separately for shallow (quaternary) and deep aquifers.*

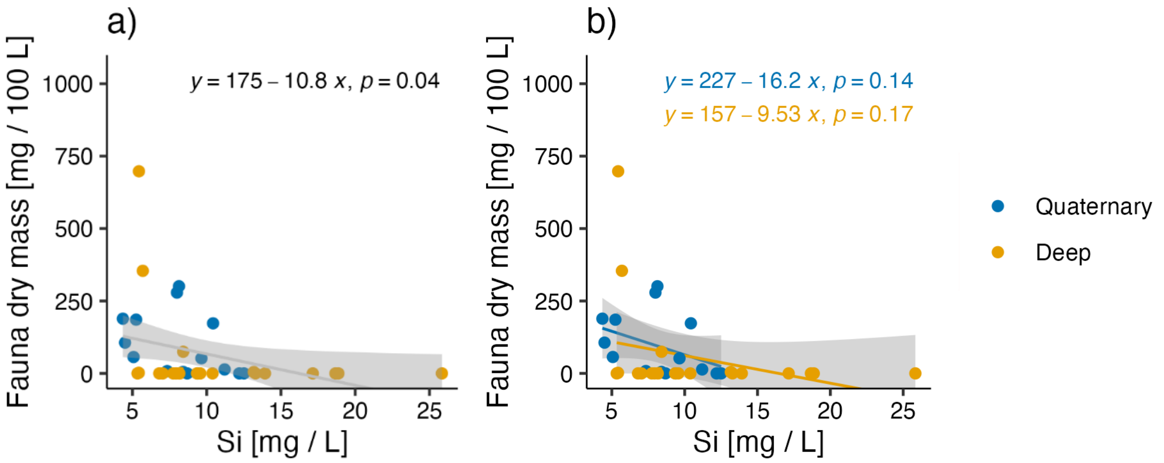

*Figure S19: Organisms in dependence on silica concentrations. Fauna biomasses a): total trend; b): Trends separate for deep and shallow (quaternary) aquifers: fauna biomass. The equations for the trends are given below the respective plot. For fauna individuals and prokaryotes, see Figure 3 in the main paper.*

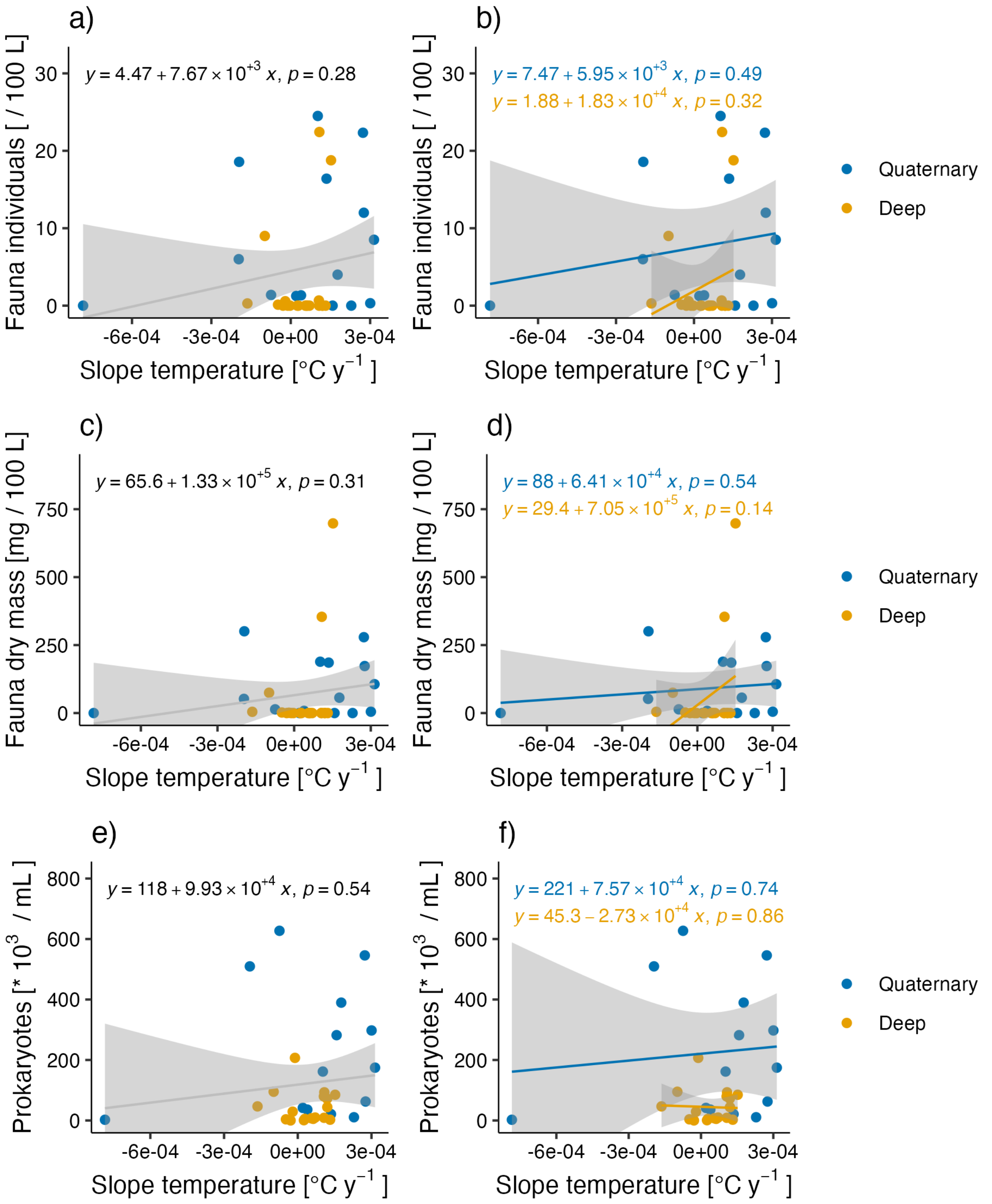

*Figure S20: Dependence of fauna individuals (top row), biomass (middle row), and prokaryotes (bottom row) on the slope in temperature over time. Same legend as in Figure 7. In b), d), and f), linear models are shown separately for shallow (quaternary) and deep aquifers.*

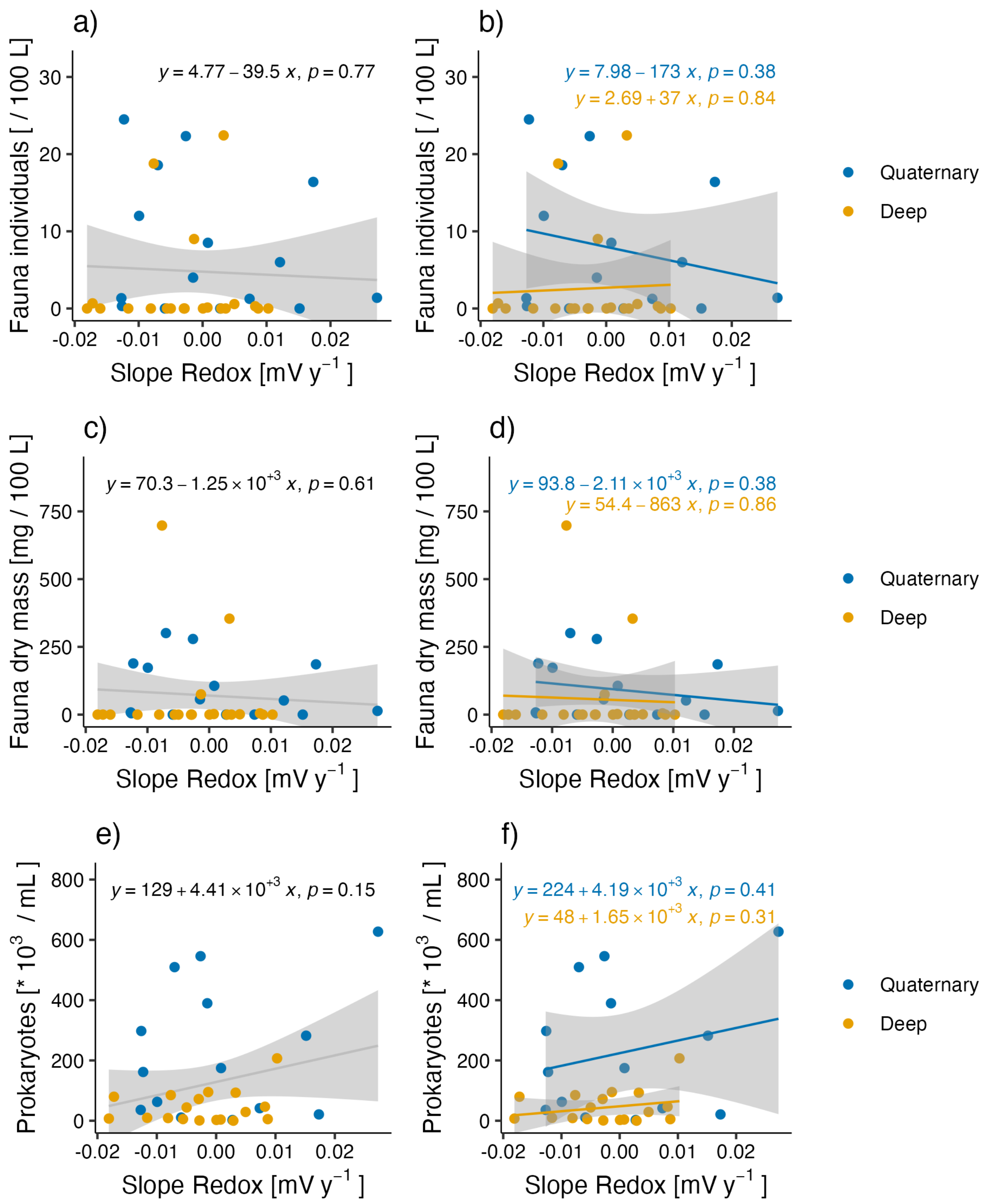

*Figure S21: Dependence of fauna individuals (top row), biomass (middle row), and prokaryotes (bottom row) on the slope in redox potential, measured in mV, yearly change. In b), d), and f), linear models are shown separately for shallow (quaternary) and deep aquifers.*

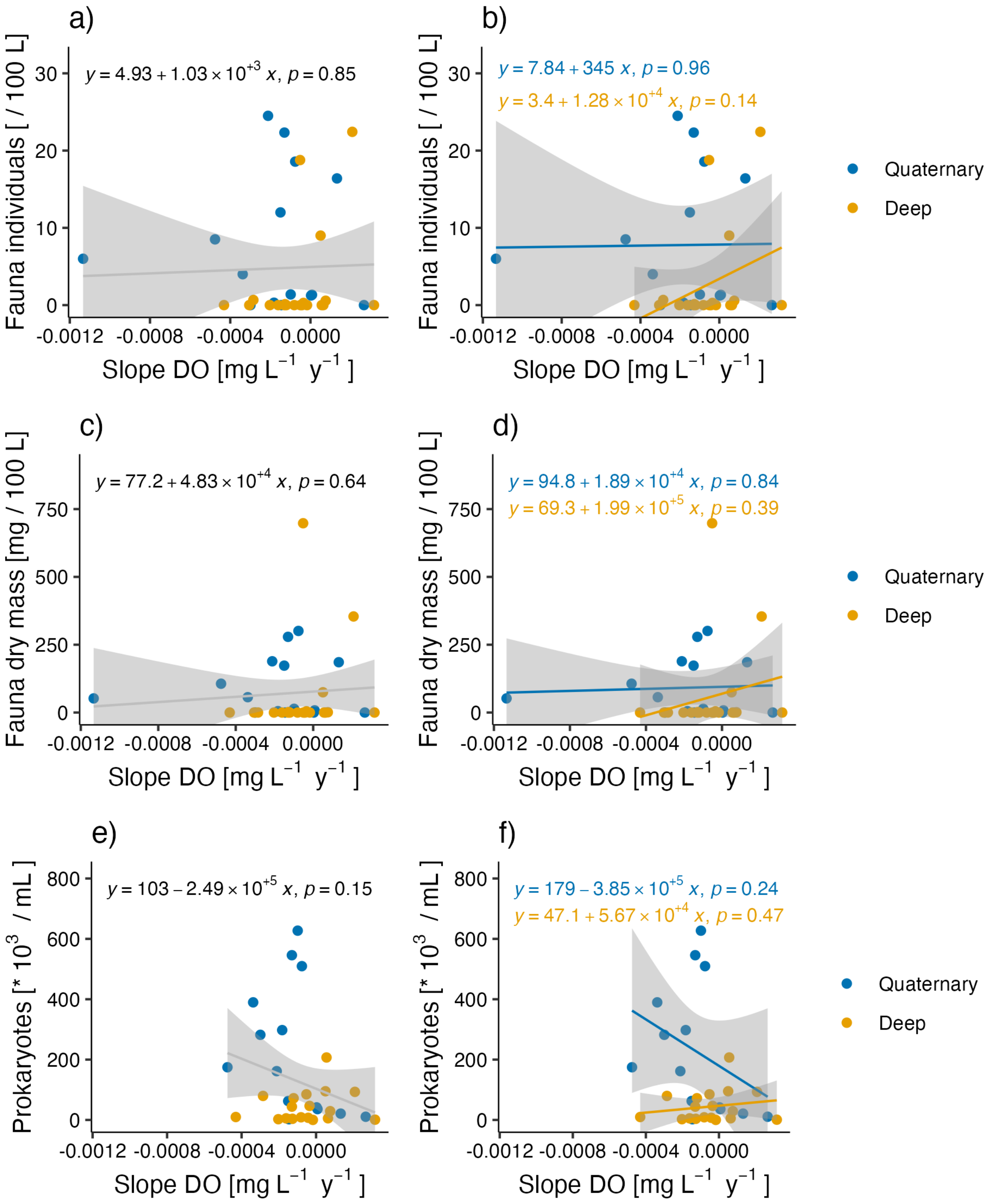

*Figure S22: Dependence of fauna individuals (top row), biomass (middle row), and prokaryotes (bottom row) on the slope in dissolved oxygen (DO), yearly change. In b), d), and f), linear models are shown separately for shallow (quaternary) and deep aquifers.*

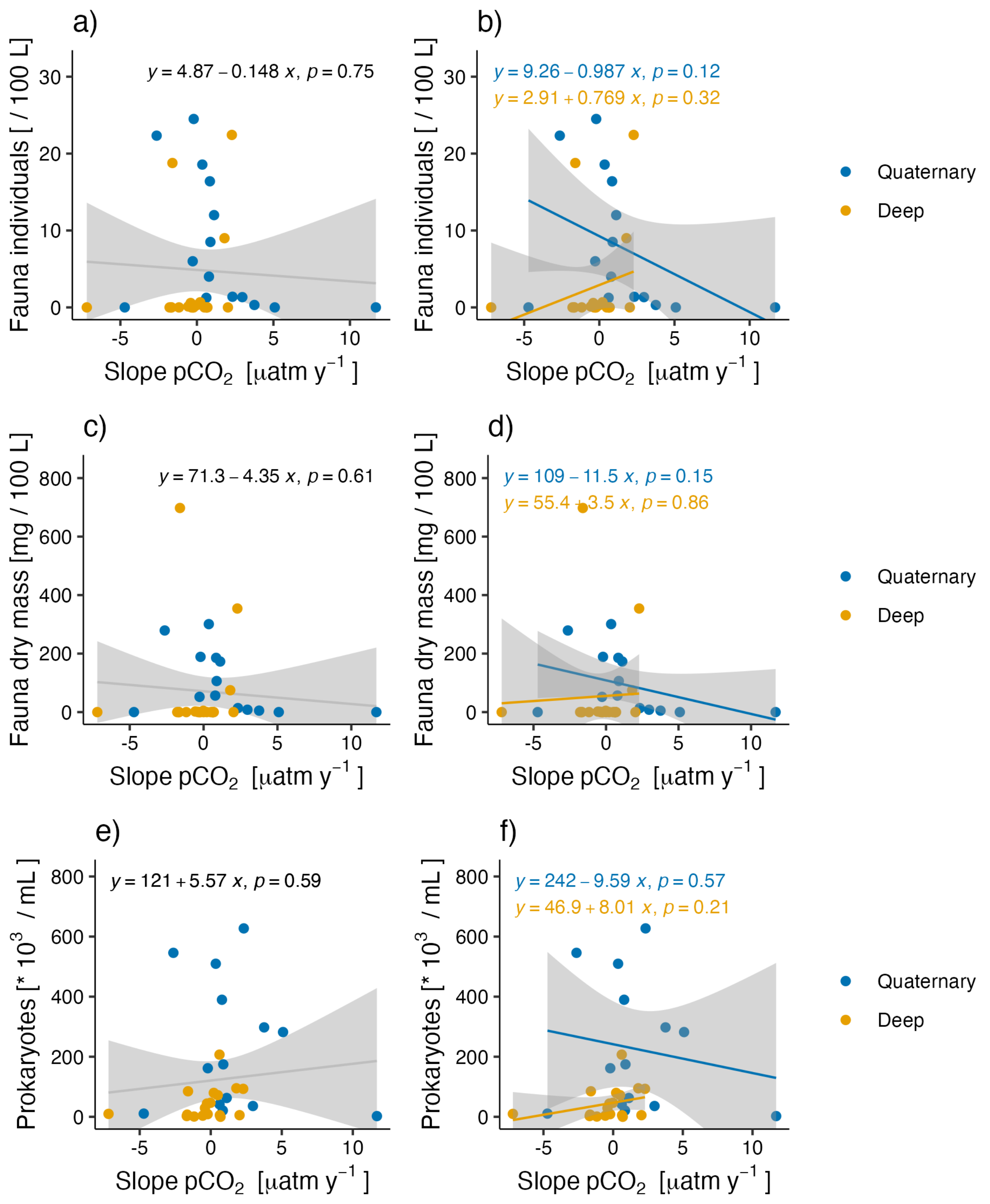

*Figure S23: Dependence of fauna individuals (top row), biomass (middle row), and prokaryotes (bottom row) on the slope in $pCO_2$), yearly change. In b), d), and f), linear models are shown separately for shallow (quaternary) and deep aquifers.*

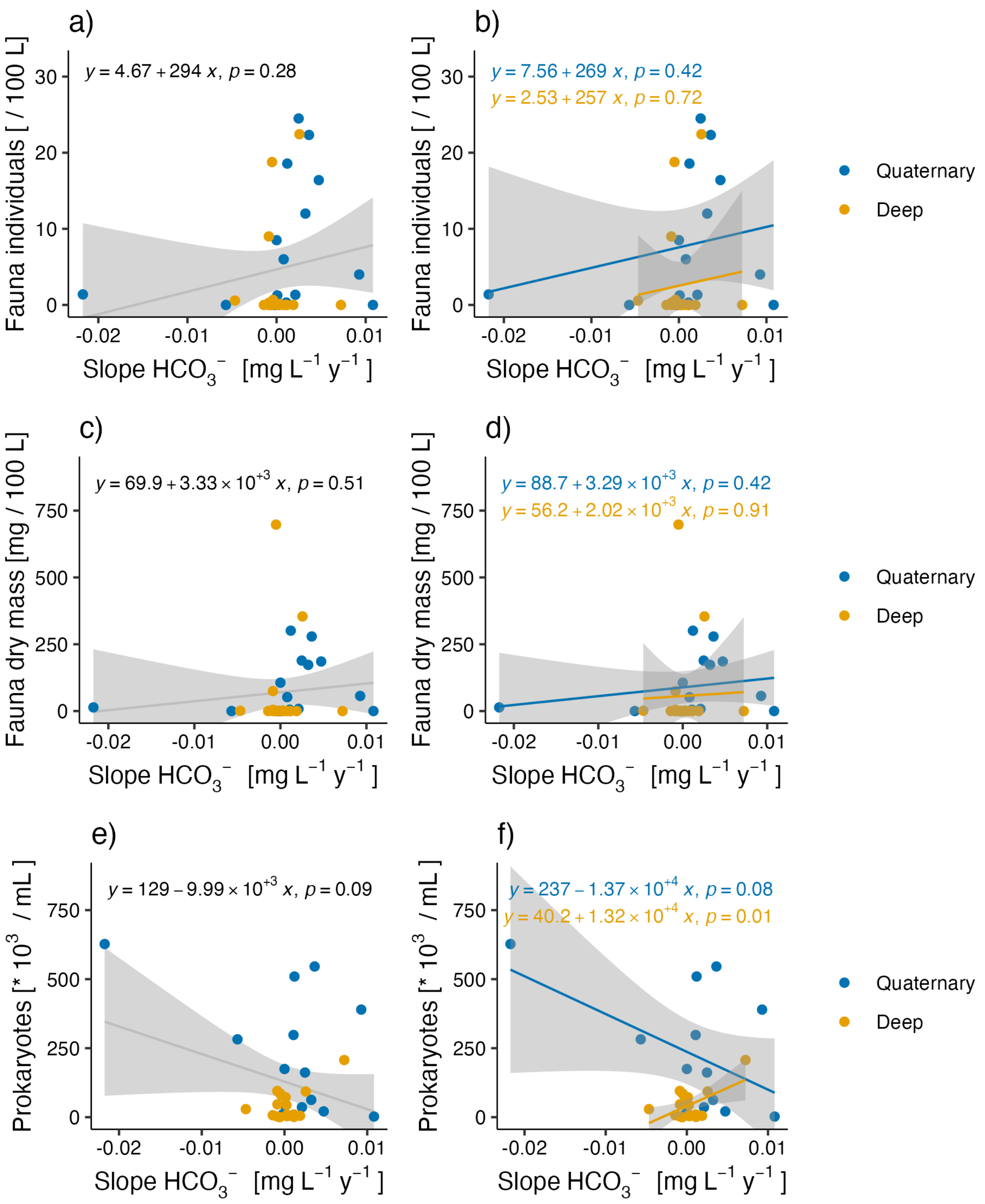

*Figure S24: Dependence of fauna biomass (top right) and prokaryotes (bottom) on the slope in HCO3 over time. Same legend as in Figure 7. For fauna individuals and prokaryotes, see the main paper.*

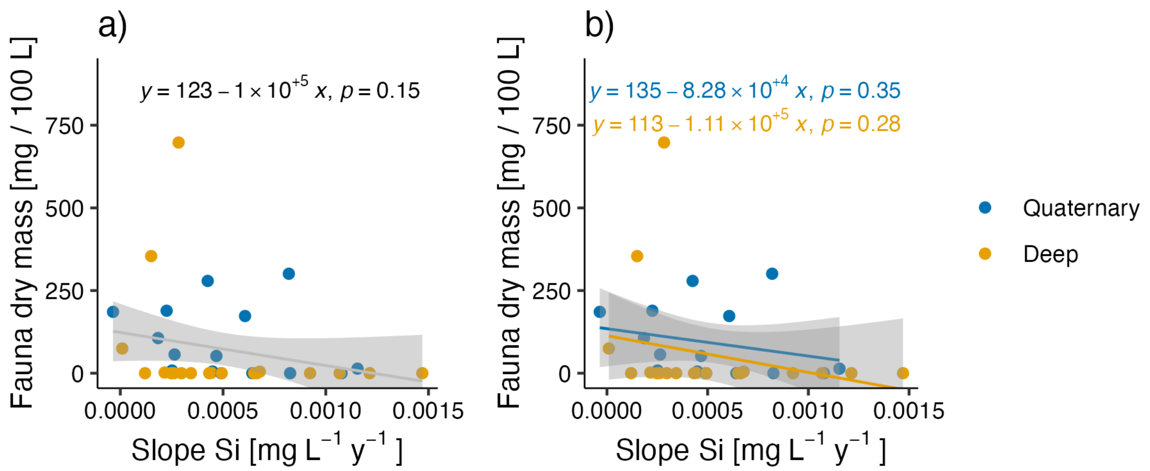

*Figure S25: Dependence of fauna biomass on the slope in silica over time. a) general linear trend; b), linear models are shown separately for shallow (quaternary) and deep aquifers. For fauna individuals and prokaryotes, see Figure 4 in the main paper.*

### Supplementary Information (SI) 10 – Results - Environmental fitting (envfit) of variables to the MDS of the faunal assemblages

*Table S4: Environmental fitting of variables to the MDS of the faunal assemblages. NMDS1 and 2 are the two NMDS axes that were used for this test. $R^2$ measure of fit; p =adjusted probability level; also indicated with stars in the last column. Chemical and physical data from 2019 to 2021, except for the standard deviations ("sd") and the slopes of the trends over time, given as "slope_per_year".*

| | NMDS1 | NMDS2 | $R^2$ | $p_{adjusted}$ | |
|---|---|---|---|---|---|
| easting | -0.44 | 0.90 | 0.01 | 0.92 | |
| northing | 0.29 | -0.96 | 0.11 | 0.18 | |
| waterlevel slope per year | -0.71 | 0.70 | 0.03 | 0.59 | |
| T (temperature, °C) | 0.17 | 0.99 | 0.15 | 0.10 | |
| EC | 1.00 | -0.05 | 0.07 | 0.39 | |
| pH | -0.03 | -1.00 | 0.01 | 0.88 | |
| Eh | 0.57 | -0.82 | 0.45 | 0.002 | ** |
| $pCO_2$ | 0.08 | 1.00 | 0.15 | 0.10 | |
| $HCO_3^-$ | 0.10 | 0.99 | 0.07 | 0.39 | |
| COD | -0.45 | 0.89 | 0.22 | 0.04 | |
| Ca | 0.94 | -0.35 | 0.20 | 0.06 | |
| Cl | 0.87 | -0.49 | 0.06 | 0.44 | |
| DO | 0.30 | -0.96 | 0.42 | 0.003 | ** |
| DOC | 0.50 | 0.86 | 0.15 | 0.10 | |
| DRP | -0.55 | 0.84 | 0.01 | 0.91 | |
| F | -0.29 | 0.96 | 0.05 | 0.49 | |
| $Fe_d$ | -0.64 | 0.77 | 0.39 | 0.00 | |
| Mg | 0.99 | 0.12 | 0.06 | 0.42 | |
| $Mn_d$ | -0.74 | 0.67 | 0.04 | 0.54 | |
| $NH_4$-N | -0.63 | 0.77 | 0.14 | 0.10 | |
| $NO_2$-N | 0.76 | 0.64 | 0.14 | 0.13 | |
| $NO_3$-N | 0.74 | -0.67 | 0.46 | 0.002 | ** |
| Na | 0.83 | 0.55 | 0.03 | 0.71 | |
| Si | -0.85 | 0.53 | 0.35 | 0.004 | ** |
| $SO_4^{2-}$ | 0.94 | -0.35 | 0.23 | 0.03 | |
| sd T | 0.93 | 0.37 | 0.46 | 0.001 | ** |
| sd $SO_4^{2-}$ | 0.99 | 0.15 | 0.39 | 0.002 | ** |
| sd COD | -0.34 | 0.94 | 0.08 | 0.29 | |
| sd Cl | 1.00 | 0.05 | 0.04 | 0.56 | |
| sd DO | 0.70 | -0.72 | 0.10 | 0.22 | |
| sd DOC | 0.11 | 0.99 | 0.05 | 0.51 | |
| sd DRP | -0.76 | 0.65 | 0.12 | 0.19 | |
| sd $NO_3$-N | 0.92 | -0.39 | 0.45 | 0.002 | ** |
| sd EC | 0.99 | -0.14 | 0.14 | 0.14 | |
| sd pH | 0.09 | 1.00 | 0.12 | 0.16 | |
| sd Si | -0.58 | 0.81 | 0.11 | 0.17 | |
| sd $pCO_2$ | -0.09 | 1.00 | 0.03 | 0.69 | |
| sd $HCO_3^-$ | 0.50 | 0.87 | 0.12 | 0.16 | |
| sd Eh | 0.56 | -0.83 | 0.18 | 0.07 | |
| COD slope per year | -0.87 | 0.50 | 0.09 | 0.24 | |
| DO slope per year | -0.73 | 0.68 | 0.04 | 0.58 | |
| DOC slope per year | -0.29 | -0.96 | 0.09 | 0.29 | |
| $SO_4^{2-}$ slope per year | -0.97 | 0.25 | 0.14 | 0.11 | |
| $NO_3^-$ slope per year | -0.92 | -0.40 | 0.03 | 0.65 | |
| DRP slope per year | 0.92 | -0.40 | 0.02 | 0.73 | |
| EC slope per year | -0.94 | 0.33 | 0.04 | 0.52 | |
| pH slope per year | 0.91 | 0.41 | 0.05 | 0.49 | |
| Si slope per year | -0.50 | 0.87 | 0.22 | 0.03 | * |
| $pCO_2$ slope per year | 0.25 | -0.97 | 0.01 | 0.82 | |
| T slope per year | 0.96 | -0.27 | 0.08 | 0.27 | |
| $HCO_3^-$ slope per year | 0.43 | -0.90 | 0.03 | 0.68 | |
| Eh slope per year | 0.18 | 0.98 | 0.11 | 0.20 | |
| prokaryotes | 1.00 | 0.07 | 0.31 | 0.01 | * |

**References**

Andrews, J.A., Schlesinger, W.H., 2001. Soil CO2 dynamics, acidification, and chemical weathering in a temperate forest with experimental CO2 enrichment. Global Biogeochemical Cycles 15, 149–162. https://doi.org/10.1029/2000GB001278

Bailey, R.A., 1991. Chemistry of the Environment.

Bakalowicz, M., 2005. Karst groundwater: A challenge for new resources. Hydrogeology Journal 13, 148–160. https://doi.org/10.1007/s10040-004-0402-9

Bakalowicz, M., 1994. 4 Water geochemistry: water quality and dynamics, in: Gibert, J., Danielopol, D.L., Stanford, J.A. (Eds.), Groundwater Ecology. San Diego, CA., pp. 97–127. https://doi.org/10.1016/B978-0-08-050762-0.50011-5

Beaulieu, E., Goddéris, Y., Donnadieu, Y., Labat, D., Roelandt, C., 2012. High sensitivity of the continental-weathering carbon dioxide sink to future climate change. Nature Clim Change 2, 346–349. https://doi.org/10.1038/nclimate1419

Blume, H.-P., Brümmer, G.W., Horn, R., Kandeler, E., Kögel-Knabner, I., Kretzschmar, R., Stahr, K., Wilke, B.-M., 2010. Scheffer/Schachtschabel Lehrbuch der Bodenkunde.

Boulton, A.J., Dole-Olivier, M.J., Marmonier, P., 2004. Effects of sample volume and taxonomic resolution on assessment of hyporheic assemblage composition sampled using a Bou-Rouch pump. Arch. Hydrobiol. 159, 327–355. https://doi.org/10.1127/0003-9136/2004/0159-0327

Brantley, S.L., Shaughnessy, A., Lebedeva, M.I., Balashov, V.N., 2023. How temperature-dependent silicate weathering acts as Earth's geological thermostat. Science 379, 382–389. https://doi.org/10.1126/science.add2922

Calmels, D., Galy, A., Hovius, N., Bickle, M., West, A.J., Chen, M.-C., Chapman, H., 2011. Contribution of deep groundwater to the weathering budget in a rapidly eroding mountain belt, Taiwan. Earth and Planetary Science Letters 303, 48–58. https://doi.org/10.1016/j.epsl.2010.12.032

Dessert, C., Dupré, B., Gaillardet, J., François, L.M., Allègre, C.J., 2003. Basalt weathering laws and the impact of basalt weathering on the global carbon cycle. Chemical Geology, Controls on Chemical Weathering 202, 257–273. https://doi.org/10.1016/j.chemgeo.2002.10.001

Fitts, C.R., 2012. Groundwater Science, Second Edi. ed. Elsevier, Amsterdam.

Gaillardet, J., Dupré, B., Louvat, P., Allègre, C.J., 1999. Global silicate weathering and CO2 consumption rates deduced from the chemistry of large rivers. Chemical Geology 159, 3–30. https://doi.org/10.1016/S0009-2541(99)00031-5

Gattuso, J., Epitalon, J., Lavigne, H., Orr, J., 2022. seacarb: Seawater Carbonate Chemistry. R package version 3.3.1.

Gislason, S.R., Oelkers, E.H., Eiriksdottir, E.S., Kardjilov, M.I., Gisladottir, G., Sigfusson, B., Snorrason, A., Elefsen, S., Hardardottir, J., Torssander, P., Oskarsson, N., 2009. Direct evidence of the feedback between climate and weathering. Earth and Planetary Science Letters 277, 213–222. https://doi.org/10.1016/j.epsl.2008.10.018

Hahn, H.J., 2003. Eignen sich Fallen zur repräsentativen Erfassung aquatischer Meiofauna im hyporheischen Interstitial und im Grundwasser? Limnologica 33, 138–146.

Hartmann, J., Jansen, N., Dürr, H.H., Kempe, S., Köhler, P., 2009. Global CO2-consumption by chemical weathering: What is the contribution of highly active weathering regions? Global and Planetary Change 69, 185–194. https://doi.org/10.1016/j.gloplacha.2009.07.007

IPCC, 2022. Climate Change 2022: Impacts, Adaptation and Vulnerability. Contribution of Working Group II to the Sixth Assessment Report of the Intergovernmental Panel on Climate Change [H.-O. Pörtner, D.C. Roberts, M. Tignor, E.S. Poloczanska, K. Mintenbeck, A. Alegría, M. Craig, S. Langsdorf, S. Löschke, V. Möller, A. Okem, B. Rama (eds.)]. Cambridge University Press. Cambridge University Press, Cambridge, UK and New York, NY, USA 1–3056. https://doi.org/10.1017/9781009325844

Karlović, I., Marković, T., Šparica Miko, M., Maldini, K., 2021. Geochemical Characteristics of Alluvial Aquifer in the Varaždin Region. Water 13, 1508. https://doi.org/10.3390/w13111508

Kasting, J.F., 2019. The Goldilocks Planet? How Silicate Weathering Maintains Earth "Just Right." Elements 15, 235–240. https://doi.org/10.2138/gselements.15.4.235

Kopáček, J., Hezlar, J., 2021. Voda na Zemi. Jihočeská univerzita v Českých Budějovicích.

Legler, C., Breitig, G., Steppuhn, G., Vobach, V. (Eds.), 1986. Ausgewählte Methoden der Wasseruntersuchung - Band I. Chemische, physikalisch-chemische, physikalische und elektro-chemische Methoden. VEB Gustav Fischer Verlag, Jena.

Lehmann, N., Stacke, T., Lehmann, S., Lantuit, H., Gosse, J., Mears, C., Hartmann, J., Thomas, H., 2023. Alkalinity responses to climate warming destabilise the Earth's thermostat. Nat Commun 14, 1648. https://doi.org/10.1038/s41467-023-37165-w

Li, Gaojun, Hartmann, J., Derry, L.A., West, A.J., You, C.-F., Long, X., Zhan, T., Li, L., Li, Gen, Qiu, W., Li, T., Liu, L., Chen, Y., Ji, J., Zhao, L., Chen, J., 2016. Temperature dependence of basalt weathering. Earth and Planetary Science Letters 443, 59–69. https://doi.org/10.1016/j.epsl.2016.03.015

Li, L., Sullivan, P.L., Benettin, P., Cirpka, O.A., Bishop, K., Brantley, S.L., Knapp, J.L.A., van Meerveld, I., Rinaldo, A., Seibert, J., Wen, H., Kirchner, J.W., 2021. Toward catchment hydro-biogeochemical theories. WIREs Water 8, e1495. https://doi.org/10.1002/wat2.1495

Luijendijk, E., Gleeson, T., Moosdorf, N., 2020. Fresh groundwater discharge insignificant for the world's oceans but important for coastal ecosystems. Nat Commun 11, 1260. https://doi.org/10.1038/s41467-020-15064-8

Macpherson, G.L., 2009. CO2 distribution in groundwater and the impact of groundwater extraction on the global C cycle. Chemical Geology 264, 328–336. https://doi.org/10.1016/j.chemgeo.2009.03.018

Macpherson, G.L., Roberts, J.A., Blair, J.M., Townsend, M.A., Fowle, D.A., Beisner, K.R., 2008. Increasing shallow groundwater CO2 and limestone weathering, Konza Prairie, USA. Geochimica et Cosmochimica Acta 72, 5581–5599. https://doi.org/10.1016/j.gca.2008.09.004

Marx, A., Dusek, J., Jankovec, J., Sanda, M., Vogel, T., van Geldern, R., Hartmann, J., Barth, J. a. C., 2017. A review of CO2 and associated carbon dynamics in headwater streams: A global perspective. Reviews of Geophysics 55, 560–585. https://doi.org/10.1002/2016RG000547

Marxsen, J., Rütz, N.K., Schmidt, S.I., 2021. Organic carbon and nutrients drive prokaryote and metazoan communities in a floodplain aquifer. Basic and Applied Ecology 51, 43–58. https://doi.org/10.1016/j.baae.2020.12.006

Matzke, D., Hahn, H.J., 2002. Vergleich der Grundwasserfauna in Lockergesteins- und in Kluftgrundwasserleitern unter vergleichender Anwendung unterschiedlicher Sammeltechniken. Abschlussbericht Projekt Az HA 3214 / 1-1.

Meybeck, M., 1993. Riverine transport of atmospheric carbon: sources, global typology and budget. Water, Air, and Soil Pollution 70, 443–463.

Moosdorf, N., Hartmann, J., Lauerwald, R., Hagedorn, B., Kempe, S., 2011. Atmospheric CO2 consumption by chemical weathering in North America. Geochimica et Cosmochimica Acta 75, 7829–7854. https://doi.org/10.1016/j.gca.2011.10.007

Négrel, P., Allègre, C.J., Dupré, B., Lewin, E., 1993. Erosion sources determined by inversion of major and trace element ratios and strontium isotopic ratios in river water: The Congo Basin case. Earth and Planetary Science Letters 120, 59–76. https://doi.org/10.1016/0012-821X(93)90023-3

Oksanen, J., Simpson, G., Blanchet, F., Kindt, R., Legendre, P., Minchin, P., O’Hara, R., Solymos, P., Stevens, M., Szoecs, E., Wagner, H., Barbour, M., Bedward, M., Bolker, B., Borcard, D., Carvalho, G., Chirico, M., De Caceres, M., Durand, S., Evangelista, H., FitzJohn, R., Friendly, M., Furneaux, B., Hannigan, G., Hill, M., Lahti, L., McGlinn, D., Ouellette, M., Ribeiro Cunha, E., Smith, T., Stier, A., Ter Braak, C., Weedon, J., 2022. vegan: Community Ecology Package. R package version 2.6-2.

Oliva, P., Viers, J., Dupré, B., 2003. Chemical weathering in granitic environments. Chemical Geology, Controls on Chemical Weathering 202, 225–256. https://doi.org/10.1016/j.chemgeo.2002.08.001

Penman, D.E., Caves Rugenstein, J.K., Ibarra, D.E., Winnick, M.J., 2020. Silicate weathering as a feedback and forcing in Earth’s climate and carbon cycle. Earth-Science Reviews 209, 103298. https://doi.org/10.1016/j.earscirev.2020.103298

Schmidt, S.I., Cuthbert, M.O., Schwientek, M., 2017. Towards an integrated understanding of how micro scale processes shape groundwater ecosystem functions. Science of the Total Environment 592, 215–227. https://doi.org/10.1016/j.scitotenv.2017.03.047

Schmidt, S.I., Hahn, H.J., Watson, G.D., Woodbury, R.J., Hatton, T.J., 2004. Sampling fauna in stream sediments as well as groundwater using one net sampler. Acta Hydrochimica et Hydrobiologica 32, 131−137. https://doi.org/10.1002/aheh.200300522

Shaughnessy, A.R., Gu, X., Wen, T., Brantley, S.L., 2021. Machine learning deciphers CO<sub>2</sub> sequestration and subsurface flowpaths from stream chemistry. Hydrol. Earth Syst. Sci. 25, 3397–3409. https://doi.org/10.5194/hess-25-3397-2021

Vinnarasi, F., Srinivasamoorthy, K., Saravanan, K., Gopinath, S., Prakash, R., Ponnumani, G., Babu, C., 2021. Chemical weathering and atmospheric carbon dioxide (CO2) consumption in Shanmuganadhi, South India: evidences from groundwater geochemistry. Environ Geochem Health 43, 771–790. https://doi.org/10.1007/s10653-020-00540-3

Walker, J.C.G., Hays, P.B., Kasting, J.F., 1981. A negative feedback mechanism for the long-term stabilization of Earth’s surface temperature. J. Geophys. Res. 86, 9776. https://doi.org/10.1029/JC086iC10p09776

Webb, B.W., Waling, D.E., 1992. 4. Water quality II. Chemical characteristics, in: Calow, Peter & Geoffrey E. Petts (Eds.): The Rivers Handbook - Hydrological and Ecological Principles. Volume One. Blackwell Science Ltd, pp. 73–100.

Weiss, R.F., 1974. Carbon dioxide in water and seawater: the solubility of a non-ideal gas. Marine Chemistry 2, 203–215. https://doi.org/10.1016/0304-4203(74)90015-2

White, A.F., Blum, A.E., Bullen, T.D., Vivit, D.V., Schulz, M., Fitzpatrick, J., 1999. The effect of temperature on experimental and natural chemical weathering rates of granitoid rocks. Geochimica et Cosmochimica Acta. https://doi.org/10.1016/S0016-7037(99)00250-1

Xiong, L., Bai, X., Zhao, C., Li, Y., Tan, Q., Luo, G., Wu, L., Chen, F., Li, C., Ran, C., Xi, H., Luo, X., Chen, H., Zhang, S., Liu, M., Gong, S., Xiao, B., Du, C., Song, F., 2022. High-Resolution

Data Sets for Global Carbonate and Silicate Rock Weathering Carbon Sinks and Their Change Trends. Earth's Future 10, e2022EF002746. https://doi.org/10.1029/2022EF002746
Zhang, S., Bai, X., Zhao, C., Tan, Q., Luo, G., Wang, J., Li, Q., Wu, L., Chen, F., Li, C., Deng, Y., Yang, Y., Xi, H., 2021. Global CO2 consumption by silicate rock chemical weathering: Its past and future. Earth's Future 9, e2020EF001938. https://doi.org/10.1029/2020EF001938
Zhang, S., Planavsky, N.J., 2019. Revisiting groundwater carbon fluxes to the ocean with implications for the carbon cycle. Geology 48, 67–71. https://doi.org/10.1130/G46408.1
